# *Drosophila* maintain a consistent navigational goal angle for days to weeks

**DOI:** 10.64898/2025.12.09.693277

**Authors:** Jazz L. Weisman, Thomas L. Mohren, James D. Ryu, Maya Z. Wyse, Eduardo Dias-Ferreira, Gaby Maimon

## Abstract

Past work has demonstrated that *Drosophila* can keep to a consistent navigational bearing for minutes to hours. Here, we ask whether they can do so over days to weeks. First, we describe an experimental rig that allows individual head-fixed *Drosophila* to live for at least two weeks within a virtual-reality environment. Flies walk on a spherical treadmill and receive sugar drops at defined moments as food. Individuals express robust circadian and sleep rhythms on these rigs. We further show that flies freely navigating an environment containing a single visual orienting cue (akin to the sun) will often pick a unique direction and walk forward along that direction for tens to hundreds of meters over days to weeks. This preferred direction can be considered a goal angle because individuals will repeatedly correct for experimentally induced virtual rotations away from this angle. Flies rely on the visual cue to effectively progress forward along the goal angle—walking in circles without it—and they return to walking forward along the same angle in the morning after spending a full night (twelve hours) in darkness without the cue. These results argue for the existence of navigation goals in the *Drosophila* brain with a persistence time of days to weeks. Furthermore, the technology introduced here may enable trained behaviors across thousands of reinforcement trials in *Drosophila*, a paradigm central to mammalian neuroscience yet absent in flies.

## Introduction

Insects, like Monarch butterflies, locusts, and some moths, famously migrate vast distances (Kennedy 1951; Chapman et al. 2015; Reppert et al. 2016; Dreyer et al. 2018; Menz et al. 2022). Less well appreciated is that even *Drosophila*, under appropriate conditions, can travel surprisingly far. For example, when tens of thousands of *Drosophila* spp. were released in a barren desert, a few individual flies were caught ∼15 km away (Coyne et al. 1982; Dickinson 2014). A more recent variant of this experiment—in which cameras captured the arrival times of flies to food traps 1 km away from a desert release site—revealed that dispersing *Drosophila* often fly straight under these conditions and do so at an average ground speed of 1 m/s (Leitch et al. 2021). When flying at 1 m/s, reaching a trap 1 km away takes approximately 17 min and reaching a trap 15 km away takes approximately 4 hours. These field experiments thus argue that *Drosophila* can maintain a consistent navigational angle for tens of minutes to hours.

In the lab, *Drosophila* can similarly maintain a consistent navigational angle for timescales of minutes to hours (Giraldo et al. 2018; Green et al. 2019; Haberkern et al. 2022; Mussells Pires et al. 2024; Westeinde et al. 2024). For example, head-fixed, walking *Drosophila* will often progress forward along a consistent direction within a visual virtual environment, especially when they are made hot and/or hungry and thus highly motivated to locomote (perhaps like the flies released in the desert, mentioned above) (Green et al. 2019; Haberkern et al. 2022; Mussells Pires et al. 2024; Westeinde et al. 2024). This behavior, called menotaxis, does not reflect the fly walking forward regardless of her direction in the virtual world, because when the fly is virtually rotated within the environment she consistently turns back to her previous orientation before resuming her forward locomotion (Green et al. 2019).

Navigational studies on *Drosophila* to date— including the study of menotaxis, but also other tasks—have focused on behaviors that flies exhibit on the timescale of seconds to hours (Heisenberg and Wolf 1984; Giraldo et al. 2018; Weir and Dickinson 2012; Green et al. 2019; Haberkern et al. 2022; Westeinde et al. 2024; Mussells Pires et al. 2024; Siliciano et al. 2025; Kim and Dickinson 2017; Corfas et al. 2019; Behbahani et al. 2021; Bahl et al. 2013; and see Götz 1987, for an example of a multiday study). The lifespan of *Drosophila*, however, is 2– 3 months. We thus wondered whether *Drosophila* might be able to express a persistent navigational behavior over longer periods. To answer this question, we built a set of easily clonable visual virtual-reality rigs, each of which can house a single head-fixed fly that remains alive for a week or more. Although technologies that allow for imaging neural activity in head-fixed *Drosophila* over multiple days have been described (Huang et al. 2018; Flores-Valle et al. 2022; Flores-Valle et al. 2025) those existing setups were optimized for studying one fly at a time in the context of fluorescence imaging rather than many flies in parallel performing a task. As a result, no coherent navigational behavior that becomes evident over days to weeks has been described in *Drosophila*. Using the new rigs we have developed, we show that individual *Drosophila* can maintain a consistent navigational goal angle for days to weeks. The angle each fly keeps can be considered an actively maintained goal direction because individuals will routinely correct for brief, experimentally induced rotations of their navigational direction. We further show that flies can return to their goal angle even after twelve hours (i.e., a full night) in darkness. These findings point to navigational signals in the *Drosophila* brain that endure for days to weeks. Given the experimental advantages of *Drosophila*, the biological substrates of such signals are now poised to be delineated.

## Results

### Experimental system

A variety of behaviors have been studied in tethered *Drosophila* (Green et al. 2019; Kohatsu et al. 2011; Sten et al. 2025; Bahl et al. 2013; Götz 1987; Tammero and Dickinson 2002; Heisenberg and Wolf 1984). Tethered paradigms are particularly important in flies because they allow one to make concurrent neurophysiological measurements during behavior. Recent advances have allowed for imaging head-fixed flies over very long periods, even days, but the rigs used for such measurements have not been optimized for motivating navigational behavior (Huang et al. 2018; Flores-Valle et al. 2022; Flores-Valle et al. 2025). Here we wished to test whether flies exhibit navigational behaviors that only become evident over multi-day timescales. As such, we designed and constructed new behavioral rigs appropriate for this task, built around the following design criteria: (1) the rigs needed to be compact, such that many could fit in a small room, controlled by a single user; (2) flies needed to be fed on the rigs, such that they could survive for days to weeks without removal; and (3) the rigs needed to be constructed in a manner that allowed for robust behavioral measurements, without mechanical or electronic failures, or excessive user intervention as experiments were running.

With these requirements in mind, we designed and constructed behavioral rigs that allowed a single head-fixed fly to survive for up to two weeks within a visual virtual-reality environment (Figure 1 A). Individual flies walked on an air-supported ball. They never needed to be removed from the rig because they received nutrition and hydration, at experimentally defined time points, from small drops of sugar solution delivered via a glass pipette (Figure 1 B). A projector was used to present visual stimuli on an immersive, conical display (Figure 1 C). This design allowed a single projector to deliver a panoramic image in a small footprint. In addition to the projector, an array of overhead white LEDs could turn on and off as a circadian cue. Each rig was compact, occupying a 24 × 16 inch footprint (Figure 1 A). The rigs were also modular and scalable, such that three fit on a 24 × 48 inch breadboard, and two sets of three can be stacked for a total of six rigs within this footprint (Figure 1 D). All the experiments described here were carried out on twelve total rigs of this sort. See Methods for more details (Figures S1, S2).

**Figure 1:**
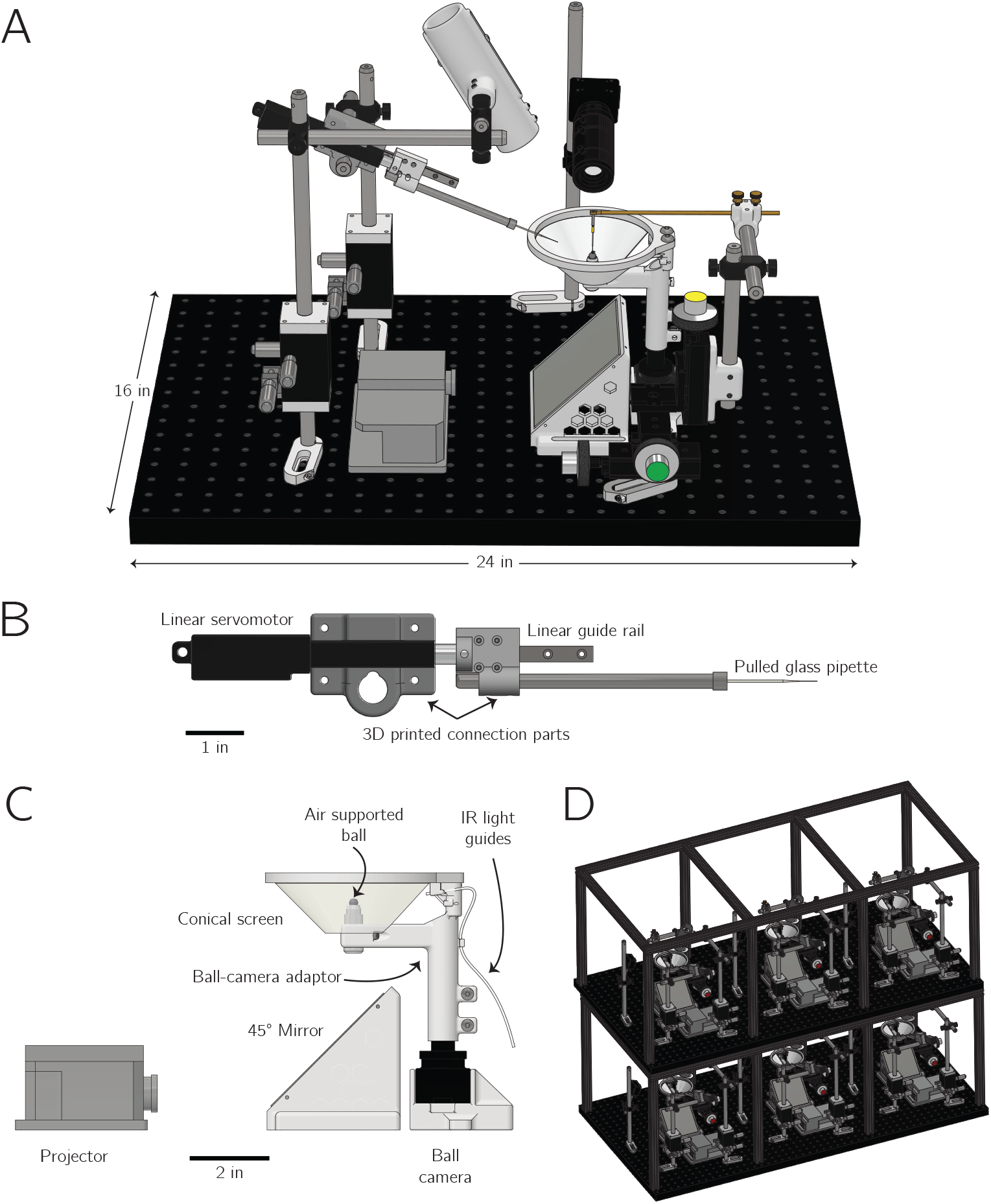
A rig for studying the navigational behavior of tethered *Drosophila* over days to weeks in parallel. **A:** Three dimensional rendering of one behavioral rig. One rig fits on a 24 × 16 inch breadboard. **B:** The feeding system, which consists of a pulled glass capillary, or pipette, that can be positioned under the fly’s proboscis with a linear servo motor. A solenoid valve regulates when a brief air-pressure pulse ejects a small sugar drop. **C:** Detail view of the projector, 45^°^ mirror, and conical screen. The fly walks on an air-cushioned ball. A camera tracks the movements of the ball from behind. **D:** Rendering of six rigs together, as they fit on a 24 × 48 inch breadboard, in two layers of three rigs each.

### Head-fixed behavior over days to weeks in *Drosophila*

In our first experiments, we measured the navigational behavior of pin-tethered flies living for 7–10 days in virtual reality. Each fly’s visual environment consisted of a single vertical bar (11^°^ wide, 60^°^ high) that rotated in closed-loop with her yaw (left/right) turns (Figure 2 A). Because the bar remained a constant size, it simulated a distant cue whose azimuthal angle informed the fly of her orientation within the environment. Overhead lights turned on and off every 12 hours, matching the phase and timing of the light cycle of the incubators in which the flies resided prior to being placed on the rig. The closed-loop bar was always present in these first experiments, meaning that the flies could infer their orientation and navigate in reference to the bar at all times, night and day (as defined by the overhead lights).

**Figure 2:**
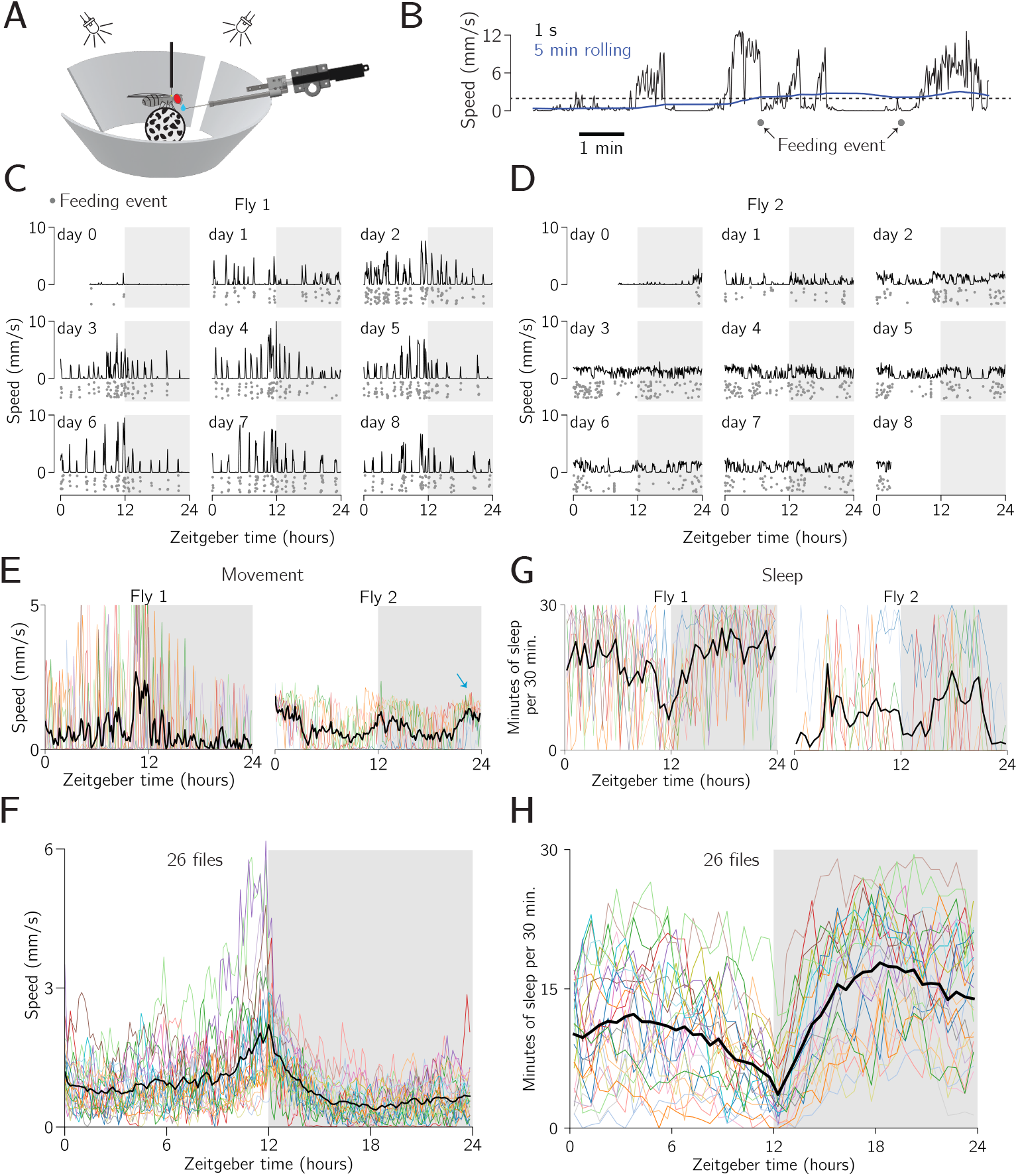
Flies walk on the ball for many days and show circadian and sleep rhythms. **A:** A fly walks on a ball with a 60^°^-tall bar that rotates in closed loop with its left/right turns. The fly is fed drops of sucrose solution multiple times per day. **B:** Example trace showing that when a 5-min rolling mean of the fly’s translation speed crosses a threshold (dotted line), the fly is fed a sugar drop. A 3-min refractory period is engaged after each feeding event constraining the shortest interval between two drops (as evident here). **C**,**D:** Mean translational speed for two example flies (5 min bins). Dots represent individual feeding events. (The *y*-positions of the feeding-event dots are jittered for clarity; all dots on a given plot happened on the same day). **E:** Mean translation speed vs. Zeitgeber time for two example flies (10 min bins). Colored lines: data from one day. Black line: mean across days. **F:** Mean translation speed vs. Zeitgeber time for the population of flies. Colored lines: data from one fly. Black line: mean across flies. **G:** Minutes of sleep per 30 minute bin vs. Zeitgeber time for the same two example flies as in panel E. Colored lines: data from one day. Black line: mean across days. **H:** Same as panel F but for sleep (i.e., 5 min+ of immobility) in each 30 minute time bin.

To promote locomotion, we made feeding events conditional on the flies walking. Specifically, to receive a sugar drop, a fly had to achieve a minimum average speed of 2 mm/s for five minutes. Additionally, there was a refractory period of three minutes after receiving a drop that constrained how soon the next drop could arrive. A pair of drop-delivery events, adherent to these rules, are depicted in Figure 2 B; the first event occurred when the 5-min rolling mean of the fly’s translation speed exceeded 2 mm/s and the second drop was delivered 3 min thereafter, that is, after the refractory period had elapsed (because the fly maintained a 2 mm/s running average for 3 min after the first drop). This feeding regimen was more successful in keeping flies alive and walking for many days compared to feeding at regular intervals. Nevertheless, many flies still did not get sufficient food with this paradigm, generally due to walking very little on the ball, and died in the first few days of the experiment. Of 81 flies initialized in this experiment, 26 survived for 7+ days (Figure S3) and were analyzed further. Future work will aim to increase the likelihood of long-term survival via technical improvements.

Some flies walked with bursts of fast walking for tens of minutes, with many long intermittent standing events (Figure 2 C; fly 1). Other flies walked relatively slowly, but more consistently over the duration of a day (Figure 2 D; fly 2). The origin of such differences in locomotor dynamics across individuals is unclear. Further example speed profiles and a summary of the speed distributions across all the flies are shown in Figure S4.

Since the flies survived for days, we wondered whether well-known long-term behavioral tendencies, like circadian rhythms and sleep (Allada et al. 2001; Patke et al. 2020; Hall 2003; Cirelli and Bushey 2008) were reproducible on these rigs. Exploring circadian rhythms, two representative flies depicted in Figure 2 C-D expressed a reliable increase in their mean walking speed in anticipation of night (*evening anticipation*) (Figure 2 E). That is, in Figure 2 E, one observes a general rise in locomotor speed in anticipation of the lights turning off (i.e., Zeitgeiber time 12). Such evening anticipation is a hallmark of an internal circadian timer because the last reliable event to which the fly could lock its behavior occurred ∼ 12 hours prior (Stoleru et al. 2004; Grima et al. 2004). Fly 2, in addition to anticipating nighttime with increased locomotion also anticipated daytime (i.e., the lights turning on at Zeitgeiber time 24 or, equivalently, 0) (*morning anticipation*, Figure 2 E, blue arrow).

Across all flies, there was a clear, statistically significant, evening anticipation peak and also a much more subtle, but also statistically significant, activity increase in anticipation of daytime (Figure 2 F). Pooling 10 min bins, the mean walking speed of the population of flies was significantly higher for the two hours before lights off (ZT 10–12) than for the middle of the daytime (ZT 4.5–7.5) (Mann-Whitney U test *p* = 4 ×10^−18^), and this was also true for the last hour of night (ZT 23–24) vs. the middle of the night (ZT 16.5–19.5) (Mann-Whitney U test *p* = 1.3 ×10^−6^). This can also be seen comparing each 10 min bin to the mid day or night group (Figure S5 B). Because morning anticipation is known to be weak or abolished in mated female *Drosophila* (Helfrich-Förster 2000; Riva et al. 2022)—which are the flies tested here— we did not expect a strong morning-anticipation peak. The clear circadian rhythms expressed by these flies argue that the circadian timer reliably entrains to the large change in overall luminance caused by the on/off state of the overhead lights. That is, the system does not get confused by the ever-present bright bar on the screen, which provides some illumination even during nighttime (except in moments when the flies choose to position the bar in the rear of their visual field, in which case the bar was not displayed due to a 40^°^ gap in our display).

In addition to circadian rhythms, sleep levels can serve as another measure of the well-being of the flies on the ball. Periods of immobility that last 5 min or longer are often used as a proxy for sleep in *Drosophila* (Huber et al. 2004; Shaw et al. 2000). During such 5 min bouts, flies have elevated thresholds for responding to sensory stimuli and the number 5 min bouts of quiescence rebound if flies are prevented from experiencing these bouts (Hendricks et al. 2000), supporting the interpretation that they represent, or correlate with, moments of actual sleep. We plotted sleep levels—operationally defined as the number of minutes of immobility, only considering 5 min or longer bouts, in each 30 min time window—for two example flies (Figure 2 G). To do this, we needed to choose a threshold speed below which the fly was considered immobile. The shape of these sleep vs. Zeitgeber time curves was robust to the exact speed threshold used, but the amount of sleep estimated in absolute terms varied as a function of this threshold. Across thresholds ranging from 0.25 mm/min (which is so low that measurement noise becomes a concern) to 60 mm/min (slow walking), average sleep levels per night varied from ∼10 to ∼21 min of sleep per 30 min (Figure S5 C).

The sleep curves shown in Figure 2 H (using a speed threshold of 1.5 mm/min) are consistent in their shape with those reported in the literature. Flies have been shown to express low levels of sleep at ZT 0 and ZT 12, with a clear anticipatory drop in average sleep before lights off (ZT 12), and, in some cases, a smaller anticipatory drop before lights on (ZT 24) (Huber et al. 2004; Shaw et al. 2000; Li et al. 2021), thus largely mirroring circadian rhythms. Flies have also been shown to express their highest levels of sleep in the middle of the night and day, with quantitatively higher values at night (Huber et al. 2004; Shaw et al. 2000; Li et al. 2021). We observed all these trends in our data, with one difference being that our flies slept less during the night than in other studies. It has been reported that flies sleep ∼570 min per night, or an average of ∼24 min per 30 min bin (Andretic and Shaw 2005). Our flies slept 330 min per night, or ∼14 min per 30 min bin, on average. While this number was sensitive to the movement threshold discussed above, the amount of sleep we registered in our apparatus was lower than published values across all of thresholds tested (Figure S5 C). Socially deprived flies have been shown to sleep fewer minutes total, and in shorter bouts, than socially satiated conspecifics (Ganguly-Fitzgerald et al. 2006; Li et al. 2021). Hungry flies also sleep less than non-hungry conspecifics (Keene et al. 2010; Sonn et al. 2018). Because our flies were both socially deprived and often hungry, these factors might have contributed to their quantitatively lower levels of sleep. Despite these differences, preparations like the one described here, and other similar ones (Huang et al. 2018; Flores-Valle et al. 2022; Flores-Valle et al. 2025), provide a potentially important new avenue for studying sleep and circadian rhythms in *Drosophila*, as they allow for the continuous tracking of behavior and posture at high spatiotemporal resolution over many days.

### Walking flies express a preferred walking direction in the virtual world across days

Beyond sleep and locomotor rhythms, we were interested in the flies’ navigational behavior on these rigs. When we plotted their trajectories, we were struck by how far many individuals displaced from their starting location across days. In Figure 3 A, we show an individual that displaced 157 m (52,000 body lengths; Fly 2) and a second that displaced 264 m (88,000 body lengths; Fly 3). The fact that flies seemed to keep a rather straight trajectory in this virtual environment reminded us of menotaxis: the behavioral tendency mentioned in the Introduction in which heated and starved flies pick a direction and walk forward along that direction for minutes to hours (Green et al. 2019). This previously described behavior, which we will refer to herein as *short-term menotaxis*, has served a key role in helping to define navigation-related computations in the fly brain (Green et al. 2019; Mussells Pires et al. 2024; Giraldo et al. 2018; Westeinde et al. 2024; Haberkern et al. 2022). In contrast, we will operationally refer to the tendency of flies to maintain a consistent navigational direction for days to weeks as *long-term menotaxis*.

**Figure 3:**
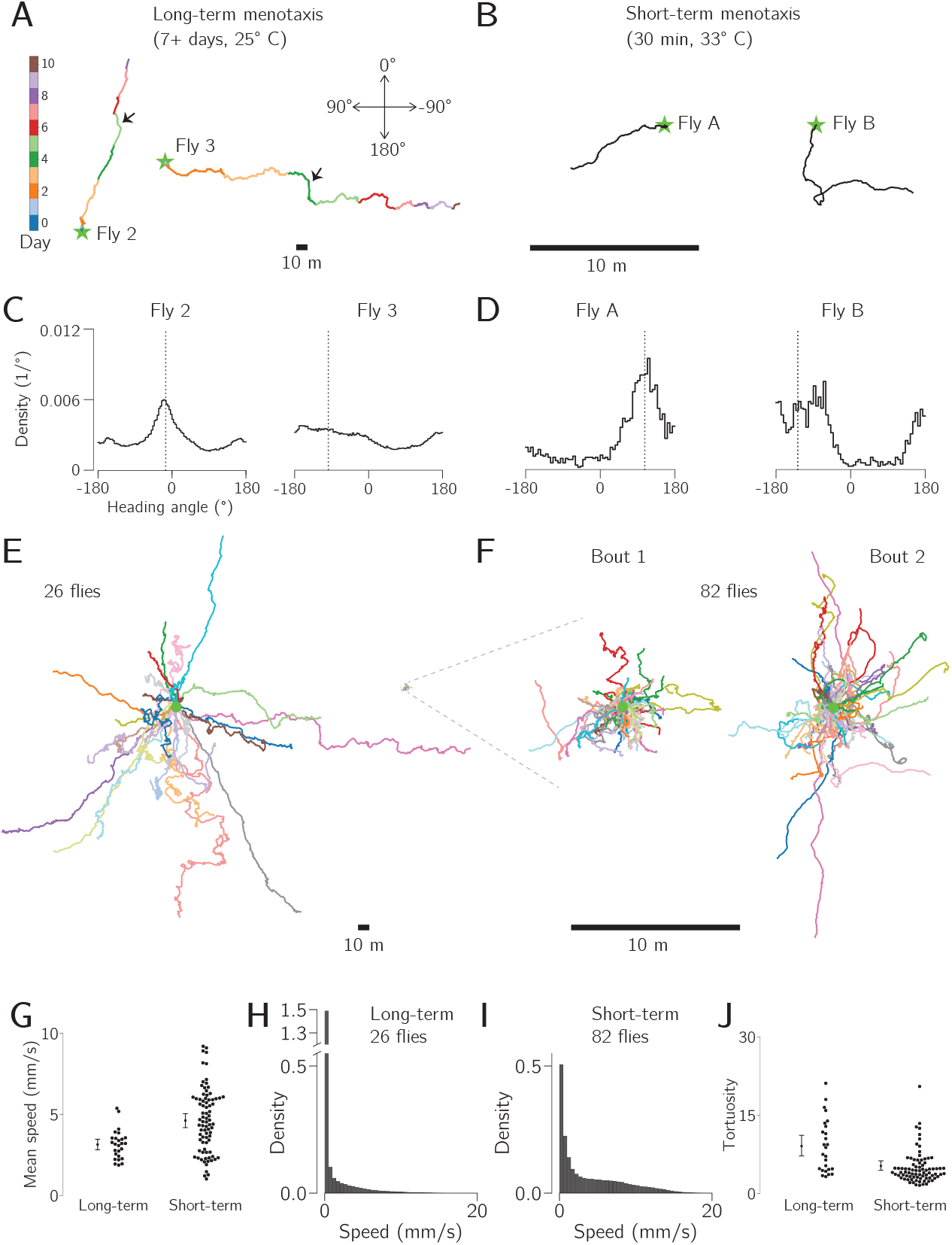
Walking flies express a preferred navigational direction over 7–10 days. **A:** Trajectories of two flies performing long-term menotaxis over 7–10 days, colored by the day of the experiment. The start location is indicated by a star, and exemplar periods of walking in a different direction from the overall average direction are indicated with black arrows. **B:** Trajectories of two example flies performing short-term menotaxis for 30 min. These flies were heated and starved (Methods). Note change in scale from panel A. **C:** Histograms of the heading angle during walking (i.e., standing events excluded) for each of the two flies from panel A. A heading angle of 0^°^ indicates walking with the bar directly in front. The angle of the vector connecting the first and last points in the trajectory on panel A is indicated by the dotted line. **D:** Same as C, but for the two short-term menotaxis flies from panel B. **E:** Trajectories of 26 flies performing long-term menotaxis over 7–10 days. Short-term menotaxis trajectories are replotted at the same spatial scale here (right). **F:** Trajectories of 82 heated and starved flies performing short-term menotaxis, 30 min trajectories. Each fly performed two bouts of short-term menotaxis and these are shown separately (see text). **G:** Average speed of flies performing long-and short-term menotaxis (standing bouts excluded). Mean and bootstrapped 95% CI shown left of each set of points. **H:** Histogram of walking speed in 1 s bins for the flies performing long-term menotaxis from panel E. Note discontinuity in the *y* axis to help show the large number of data points in the 0 – 0.5 mm/s (standing) bin while also not obscuring the shape of the distribution. **I:** Histogram of average walking speed in 1 s bins for the flies performing short-term menotaxis from panel F. **J:** Tortuosity for each fly in each data set.

To compare long-and short-term menotaxis, we collected a separate data set from heated and starved flies performing the short-term variant on the same rigs. Two representative individuals are shown in Figure 3 B. These two flies maintained a relatively narrow distribution of heading angles while walking; by comparison, the two example flies performing long-term menotaxis in Figure 3 A, exhibited broader distributions (Figure 3 C-D). Although the trajectories of the two flies performing long-term menotaxis in Figure 3 A showed a consistent preferred direction over the 7–10 day period, they both had clear periods (sometimes half a day or longer) where forward progress was oriented along a direction that deviated significantly from the overall mean direction (Figure 3 A, black arrows). The trajectories of 26 flies performing long-term menotaxis are shown in Figure 3 E and 82 flies performing short term menotaxis are shown in Figure 3 F. The short-term flies performed a pair of 30 min bouts, separated by half an hour where the bar was turned off. Each bout is shown separately in Figure 3 F. Consistent with the picture suggested by the example distributions shown in Figure 3 C-D, our population of 82 flies performing short-term menotaxis showed significantly narrower heading distributions than those of 26 flies performing long-term menotaxis (Figure S6 A, circular variance Mann-Whitney U test, *p* = 4 ×10^−8^, see Methods). The more meandering nature of the 7–10 day trajectories—further supported by the tortuosity analysis below—is one main reason that, at present, we refer to short-and long-term menotaxis as different behaviors (even if they might ultimately be shown, at least in part, to rely on similar neurons and computations). Over 7–10 days, the 26 long-term flies achieved a path-integrated distance of anywhere between a few hundred meters to *>*1 km, equivalent to 100,000–400,000 body lengths. All of the individual long-term trajectories are shown separately, colored by experimental day, in Figure S6 B.

The trajectories of flies performing short and long-term menotaxis appeared generally similar, despite differing by approximately 30-fold in spatial extent and 300-fold in temporal duration (Figure 3 E-F). However, underneath this surprising similarity, there were several important differences between the behaviors. First, flies performing short-term menotaxis walked faster in general, even when excluding standing events, than flies performing long-term menotaxis (Mann-Whitney U test, *p* = 2 ×10^−4^, Figure 3 G). Second, the long-term flies spent much more time standing still; flies performing short-term menotaxis spent ∼18% of their time with a mean speed of <0.25 mm/s (i.e., standing still), whereas flies performing long-term menotaxis spent 69% in that state (Figure 3 H-I). Finally, the long term trajectories had a higher tortuosity (i.e., the ratio of the trajectory’s path length to displacement) compared to the short-term trajectories (Figure 3 J, *p* = 4 ×10^−5^, Mann-Whitney U test).

Both the lower speed and the increased tortuosity in the trajectories of flies performing long-term menotaxis likely reflects the fact that these flies were not as motivated to walk or disperse from their current location compared to flies performing short-term menotaxis. Specifically, to motivate short-term menotaxis, we heated flies to ∼33^°^C and they were not fed for 16 to 28 hours prior to testing. Being hungry and overheated motivates flies to locomote, likely because finding food and/or a cool shelter is critical for long-term survival. The underlying motivation for flies to locomote in the long term experiment is not as clear. For the long-term data set shown in Figure 3 F continuous walking was required for sugar drops to arrive, meaning that hunger could serve as a strong motivator for walking. We therefore also collected a long-term dataset where flies were fed on a 15 min timer (independent of their walking behavior) and the overall trajectories of these long-term flies, alongside many of their locomotor metrics, resembled those of flies required to walk to get food (Figure S7). In the experiments with timed feeding, we found that flies could regularly survive for at least two weeks on our rigs, and one fly survived for 24 days, the longest time period tested to date. After 1–2 days on the ball, all long-term flies can be considered to be starved of both social interactions and protein in the diet. Future work should test whether one or both of these two factors might be critical motivators for long-range dispersal on the days to weeks timescale.

To quantify the persistence time of the longterm behavior, we transformed each day’s trajectory into a vector pointing from the first x-y location to the last x-y location on that day. We then computed the angle between this vector and the vector of the entire trajectory up until that day (Figure 4 A). In flies that exhibit a stable walking direction over days, this deviation angle would remain low over time, as it does for the example fly shown in Figure 4 A-B. Two more example flies are shown in Figure 4 C, one of which changes its traveling direction significantly on day five. When we computed this deviation angle for all 26 flies performing long-term menotaxis, we found that its magnitude remained lower than that of a shuffled distribution after the first two days (Figure 4 D). To create the shuffled curve, for each day, we took all 26 trajectory vectors from that day (one per fly) and paired each one to a randomly chosen fly’s trajectory up until that day, without replacement. We then averaged the absolute angular difference across these vector pairs for each day. We repeated this procedure 10,000 times for each day to create a distribution whose mean and 2.5%-to-97.5% range is shown in Figure 4 D (gray region).

**Figure 4:**
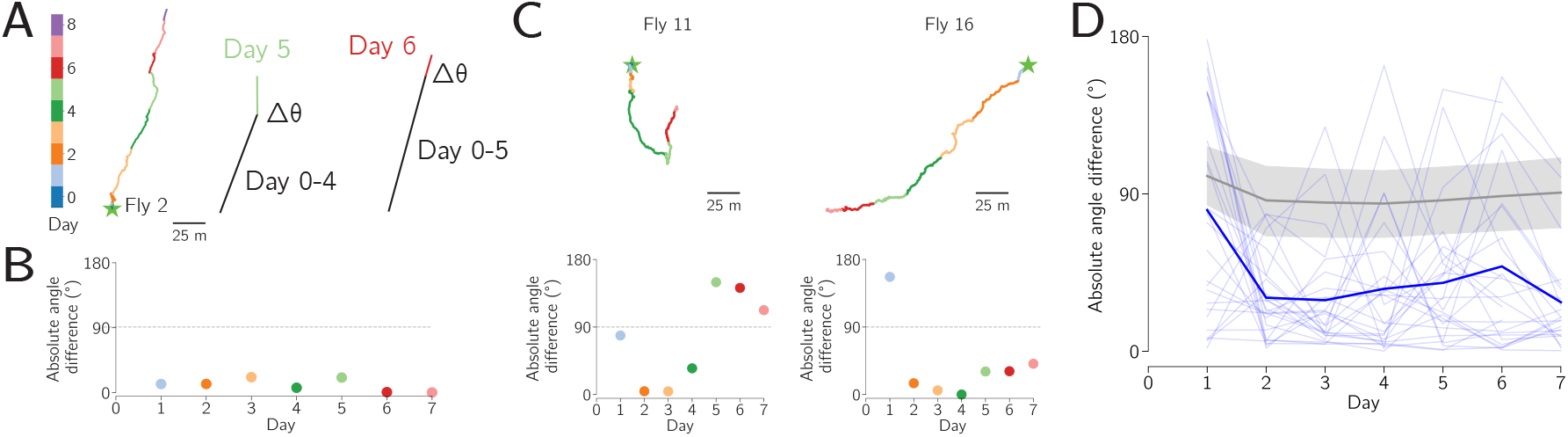
The flies’ preferred walking direction is stable over many days. **A:** Schematic of the goal stability analysis, for one example fly. Each day’s trajectory (colored line) is compared to the trajectory up to that day (black line) to compute an angle between them. **B:** Absolute angle between each day’s trajectory and the trajectory up to that day, for the example fly from panel A. Because this fly’s preferred walking direction was quite stable across days, all plotted points are near zero. **C:** Two more example flies trajectories (top) and their corresponding angle-vs-day plots (bottom). Because fly 11’s preferred walking direction changed on day 5, the plotted points became far from zero after this change in direction. Fly 11’s trajectory thus emphasizes that this analysis prioritizes detection of week-long consistency in the walking direction. **D:** Absolute angle difference for all 26 flies performing long-term menotaxis (light blue lines). Mean across flies shown in dark blue. Shuffled data and 95% confidence intervals shown in gray. Mean of actual data fell outside of the shuffled data for all points day 2 and later (*p* < 10^−4^), and for day 1, *p* = 0.0084. See Main Text for details.

### The flies’ preferred walking direction can be considered a navigational goal direction

If we make the standard assumption that our flies interpreted the closed-loop bar as a distant orienting cue (Green et al. 2019; Westeinde et al. 2024; Mussells Pires et al. 2024), then an individual walking forward with the bar at a consistent angle on the screen reflects her aiming to disperse along a consistent allocentric angle in the world (e.g., north, southeast, etc.). Progressing forward along a consistent allocentric direction for such long distances is unlikely to result from random behavior; rather, each individual likely had a specific bar position that she aimed to stabilize as she progressed forward for extended periods, with this bar position signifying a goal angle.

To test whether each fly’s preferred walking direction was an actively maintained goal angle, we rotated the flies in the virtual world 24 times per day and measured whether they corrected for these perturbations. Specifically, every hour we discontinuously jumped the closed-loop bar clockwise or counterclockwise by 90^°^. A 90^°^ bar jump mimics the fly being rotated 90^°^ in the other direction within the virtual environment. We thus refer to the event, equivalently, as a *virtual rotation* or a *bar jump*. The clockwise or counterclockwise direction of the virtual rotation was consistent for the first 24 hours of the experiment, and then it switched for the next 24 hour period, and so on, with the direction for the first day chosen randomly.

If the trajectories described in the preceding sections were the result of flies simply walking straight on the ball while ignoring the vertical bar’s position on the screen or the result of flies aiming to stabilize the bar at its current position on the screen, no matter its absolute angle, then the flies would be expected to express distorted overall trajectories in this experiment. We tested 19 flies in this virtual-rotation paradigm, for at least six days each. Two example flies exhibited an obvious preferred allocentric direction, routinely correcting for virtual rotations (Figure 5 A). Visual inspection of all the trajectories in the data set (Figure S8 A) suggested that most flies corrected for most virtual rotations.

**Figure 5:**
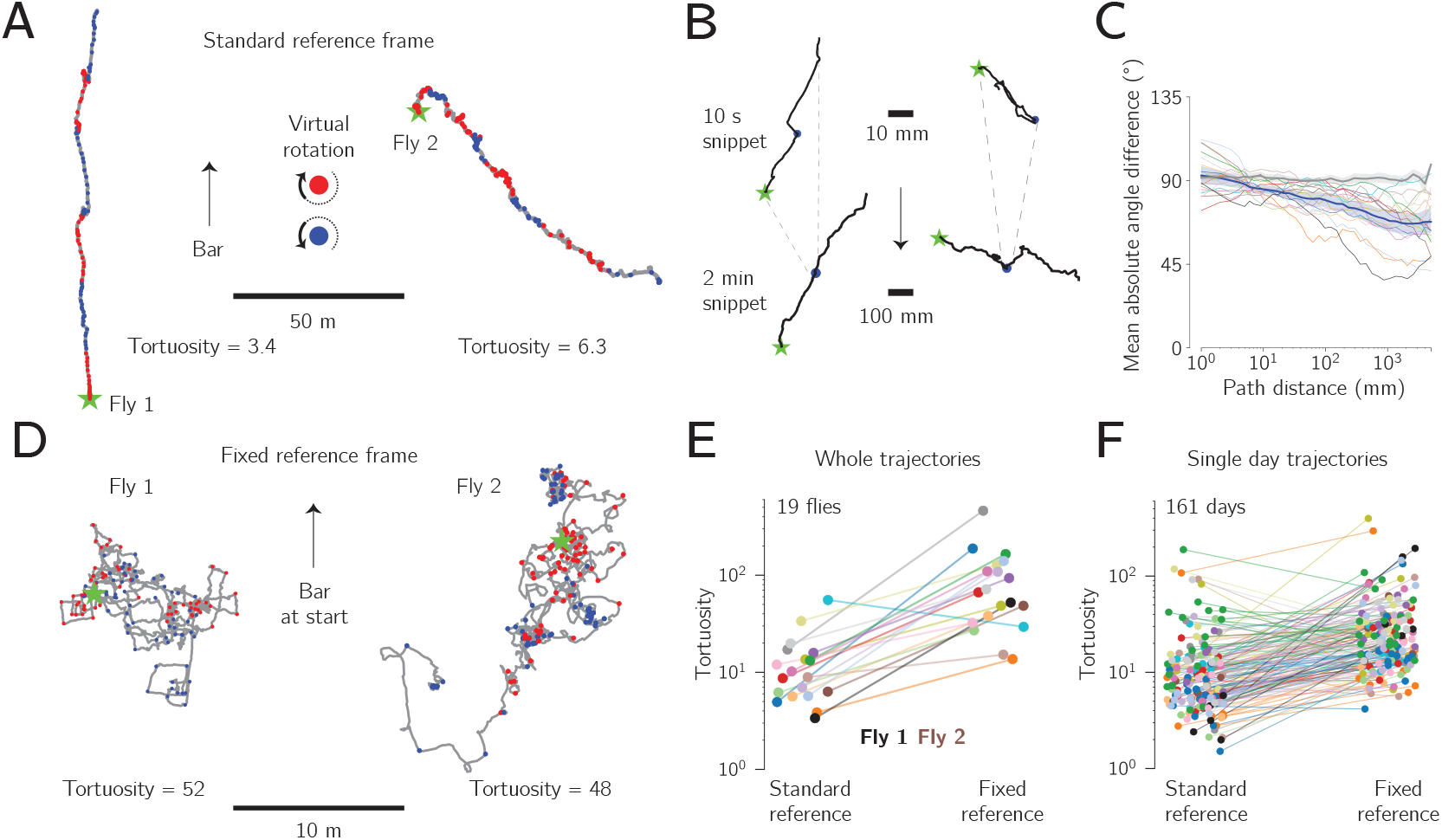
Flies have an angular goal during week-long trajectories. **A:** A pair of 7-day trajectories, where each fly experienced a 90^°^ virtual rotation (i.e., a bar jump) every hour. Red and blue dots indicate rightward and leftward virtual rotations, respectively. **B:** Top: 10 s snippet from two trajectories centered around a virtual-rotation event (blue dots). Bottom: 120 s surrounding the same virtual rotations. The longer time window reveals that the fly whose trajectory is shown on the right eventually returns to traveling along her original direction, which was not clear in the first 5 s after the rotation above. Stars indicate the start of the trajectory snippet. **C:** Mean absolute angular difference between the path-integrated vector following a bar jump and the path-integrated vector from the beginning of the trajectory until the same point (as a measure of the fly’s overall, potential, goal direction). Colored lines: individual flies. Blue line and shaded region: mean and bootstrapped 95% CI across flies. Grey line: null distribution generated from pseudo-bar jumps. See Main Text for details. **D:** Trajectories of the same example flies shown in panel A, but plotted in a fixed reference frame i.e., a reference frame that did not rotate with each bar jump. **E:** Comparison of whole-trajectory tortuosity in the two reference frames. Individual fly colors match those of the trajectories shown in Figure S8 A. Whole trajectories in the fixed reference frame are significantly more tortuous than those in the standard reference frame (*p* = 4 ×10^−5^, Wilcoxon signed-rank test). **F:** Single-day trajectories in the fixed reference frame are also significantly more tortuous than those in the standard reference frame (*p* = 3×10^−16^, Wilcoxon signed-rank test).

In Figure 5 B, we show the trajectories surrounding two virtual rotations at high spatial resolution. The trajectory on the left shows a fly that quickly reoriented herself to her preferred direction, taking only ∼1 s to correct (Figure 5 B, upper left). The trajectory on the right shows a fly that initially inverted her traveling direction after the virtual rotation, progressing forward for a few seconds along this anomalous angle (Figure 5 B, upper right trace). Examining more of her trajectory (±60 s around the bar jump), however, reveals that after the brief period of heading in an inverted direction, this fly progressed at 90^°^ to her initial angle for ∼40 s and then ultimately returned to traveling along the previous heading angle.

To quantify the flies’ behavior in response to the virtual rotations, we calculated the vector angle of the trajectory for different path-integrated distances walked after each rotation. We also calculated the angle of the fly’s overall trajectory from its start and up to the same time point (as an expression of a goal angle, if present). For each fly, we plotted the mean absolute difference between these two angles as a function of the path distance covered post rotation (Figure 5 C). The fact that the lines in Figure 5 C trend downward means that all the flies tended to turn so as to align themselves closer to the overall trajectory’s angle (i.e., the putative goal angle) after the bar jump compared to walking straight after the virtual rotation. Had the correction been complete, the lines in Figure 5 C would reach 0^°^, which they do not; this likely reflects the substantial short-scale variability of long-term menotaxis. For comparison, we created a null distribution by analyzing pseudo bar jumps in the time points between the actual bar jumps. Specifically, we rotated the trajectory after the pseudo bar jump by 90^°^ and followed the same analysis as above. When analyzing the pseudo bar jumps in the same way, flies tended to continue walking straight, on average, as expected (Figure 5 C, black line). The mean absolute angular differences for the actual bar jumps were significantly lower than the pseudo-bar-jump distribution for all path distances considered that were greater than 5 mm (Mann-Whitney U test, *p* <= 0.001, plotted in Figure S8 B).

In a second quantification of the trajectories from the virtual-rotation paradigm, we compared each fly’s standard trajectory—that is, the trajectory that uses the closed-loop bar’s angular position to assign a heading in the virtual world—with a modified trajectory in which we ignored the fact that the bar ever jumped. That is, for the modified trajectory we determined the path based on continuously tracking the ball’s rotations over time, as if the bar jumps never occurred, creating a “fixed” reference frame for the trajectory. Replotting the trajectories of the two example flies in this manner increased their tortuosity from 3.4 and 6.3 in the standard reference frame (Figure 5 A) frame to 52 and 48 in the fixed one (Figure 5 D). Indeed, the many 90^°^ turns visible in the fixed-reference frame trajectories (Figure 5 D) reflect the fact that flies were making corrective turns on the ball after each bar jump.

All but one of the fly trajectories analyzed across the 6+ days had shorter displacement and higher tortuosity values when analyzed in the fixed reference frame compared to the standard reference frame (Figure 5 E, *p* = 4 ×10^−5^, Wilcoxon signed-rank test). We interpret this result to mean that even flies that did not exhibit an obvious single goal angle across a full week nevertheless navigated by making reference to the bar on shorter timescales. We also performed the same analysis on single-day trajectories from all flies (Figure 5 F) and found that they, likewise, were more tortuous in the fixed reference frame compared to the standard reference frame (*p* = 3 ×10^−16^, Wilcoxon signed-rank test). On 85% (137 of 161) of individual days, the flies’ tortuosity was also greater in the fixed compared to the standard reference frame, as expected (upward sloping lines in Figure 5 F). Of the 15% of days were we found the opposite result, half of these came from just three flies, and 13 of the 19 flies had either zero (7 flies) or one (6 flies) day with this result.

### No evidence for time compensation in the navigational goal angle

If flies considered the closed-loop bar as a celestial object, like the sun, and they wished to head in a consistent direction in the world over a full day then they should adjust their goal angle relative to the bar in a systematic fashion as the day progresses to account for the sun’s movement across the sky. The amount of adjustment expected depends on each animal’s putative ephemeris model for how the sun progresses across the sky, which in turn depends on latitude and time of year. There is evidence that both honeybees and Monarch butterflies can compensate for the circadian movements of the sun (Lindauer 1960; Mouritsen and Frost 2002; Perez et al. 1997). Perhaps flies do likewise? We analyzed the flies’ walking trajectories and could not find supportive evidence that *Drosophila* adjust their goal angle so as to compensate for a circadian movement of the sun (Methods, Figure S9). These findings are consistent with similar experiments on *Drosophila* performing menotaxis in tethered flight, where the flies’ goal angles seemed similar across a 6-hour gap between measurements (Giraldo et al. 2018). Because there is little reason to think that *Drosophila* in the wild ever walk or fly for many hours or days with the purpose of arriving to a specific two-dimensional location, a lack of time-compensation makes sense, as has been previously argued (Giraldo et al. 2018; Warren et al. 2019). If the flies were interpreting the bar as a terrestrial cue, like a distant mountain, rather than as the sun, then we would also not expect to see evidence for time compensation.

### The navigational goal angle remains consistent after twelve hours without a visual cue

Past work has demonstrated that short-term menotaxis fundamentally relies on a neural signal in the fly brain, expressed in EPG neurons within the central complex (Green et al. 2019). EPG cells are a population of neurons whose dendrites tile a torus-shaped structure, called the ellipsoid body (Wolff et al. 2015). There is a region of locally elevated calcium activity, i.e., a “bump”, at one angular position within the ellipsoid body (Seelig and Jayaraman 2015). The bump is stable when the fly stands still. When the fly turns, the bump’s angle around the ellipsoid body (i.e., the EPG phase) changes in a corresponding manner to track the fly’s heading, akin to a rotating compass needle (Seelig and Jayaraman 2015; Green et al. 2017). For technical reasons, we have not yet been able to silence EPG cells in the context of flies attempting to perform long-term menotaxis, to directly test for a role in this behavior. We have, however, been able to perform a different experiment that has constrained the underlying mechanisms.

A remarkable aspect of EPG cells is that— analogous to mammalian head-direction cells— these neurons learn to map the angular position of relevant visual features in the external world to their population-level heading estimate (Taube et al. 1990a; Taube et al. 1990b; Seelig and Jayaraman 2015). As a result of this visual learning, the angle of the bar on the display and the angular position of the EPG bump around the ellipsoid body show a different offset from fly to fly; this offset also can change across time within a single fly (Seelig and Jayaraman 2015; Green et al. 2017). To begin to constrain the neural mechanisms of long-term menotaxis, we sought to test whether flies performing this behavior could return to walking along their previous goal angle after a very long (12 hour) period during which the closed-loop bar was invisible. If flies could do so it would suggest that either (1) the central complex has a mode in which the visual mapping to the EPG system remains stable for a notably long twelve-hour period, during which all reinforcing visual input is missing, (2) that long-term menotaxis does not rely on the EPG system in the same manner as does short-term menotaxis, or (3) that the EPG system does remap, but that downstream circuitry can somehow compensate for this in allowing the fly to maintain a consistent allocentric direction.

Our experimental paradigm was as follows (Figure 6 A). On days 1, 2, 3, 5, and 7, flies experienced the standard protocol. That is, the closed-loop bar was on 24 hours per day and the overhead lights cycled in a circadian manner: twelve hours on and twelve hours off. On days 4, 6 and 8, we turned off the closed-loop bar in synchrony with the overhead lights, at ZT 12. The bar remained off until the over-head lights turned on again twelve hours later. This protocol yielded three daytime-nighttime-daytime triplets in which the bar was off at nighttime, and three control daytime-nighttime-daytime triplets, where the bar was on throughout (white-black-white vs white-gray-white, respectively, in Figure 6 A). We refer to a full 24 hour period as a “day” and to the hours within a day where the overhead lights are on as “day-time” and the hours where the overhead lights are off as “nighttime”.

**Figure 6:**
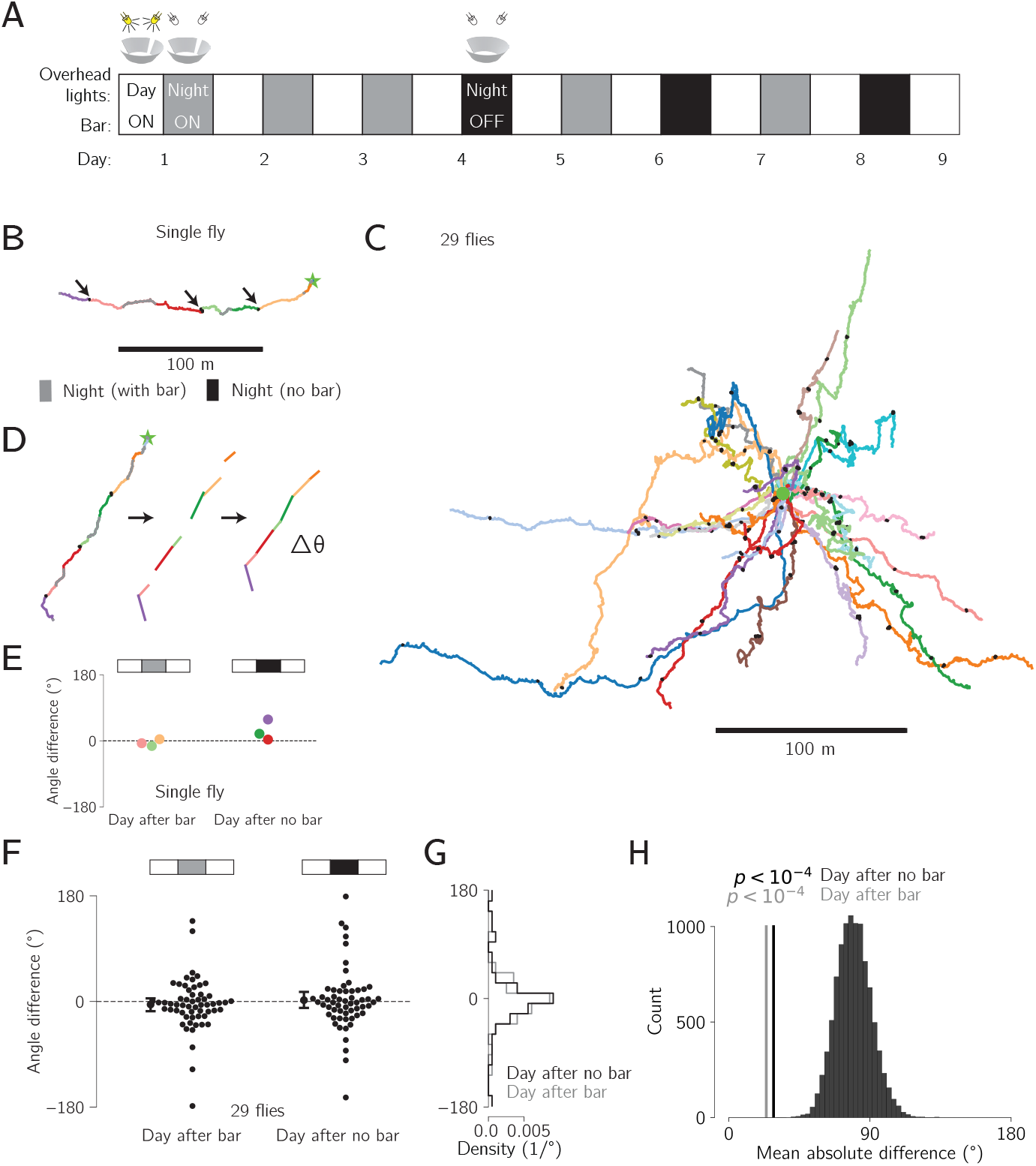
The navigational goal angle remains consistent after twelve hours without a visual cue. **A:** Experimental outline. White and grey boxes indicate the overhead lights being on and off for twelve-hour periods of daytime and nighttime, respectively. The nights when the closed-loop bar was also turned off, along with the overhead lights, are indicated with a black rather than gray box. This occurred on days 4, 6, and 8. **B:** Whole trajectory of one fly. Separate days are indicated by unique colors. Each night is colored gray when the bar was on or black when the bar was off. Arrows indicate the nights without a bar, which appear as compact points on the trajectory since the fly was not able to advance in a coherent direction in the virtual world. **C:** Trajectories of all 29 flies that had 12 hour periods without a visual cue. Each fly is indicated by a line of a different color (though some colors are shared by 2 flies). The components of each trajectory associated with nights without the bar are colored black (appearing as black points). **D:** We converted the trajectory for each day into a single vector pointing from the start point to end point of that day’s trajectory. We then calculated the angle between each successive day’s vector. **E:** Analyzing the data from the sample fly in panel D, we show the angular difference between the vector associated with each day and the previous day, colored by the second day. Days following a night with the bar are shown on the left, and days following a night without a bar are shown on right. **F:** Same as panel E but for all 29 flies. **G:** Histogram of the angular differences shown in panel D separated by type of night between the two days. **H:** Distribution of 10,000 bootstrapped mean-absolute-heading-differences between random pairs of days. Measured mean absolute difference for the day after each type of night are shown as vertical lines. *p* value, proportion of bootstrapped mean differences that are smaller than the observed mean heading difference (none for both).

Figure 6 B shows a full trajectory from an example fly. Daytime periods are indicated by a different color for each day. Nighttime periods are colored gray if the bar was on and black if the bar was off. Note that the fly did not displace significantly during bar-off nighttime periods, thus yielding black trajectories that appear as small black spots on the plot (Figure 6 B, arrows). Without a closed-loop bar, the fly cannot correct for unintended heading changes as she walks forward; these accumulating errors will produce a highly curved or meandering trajectory in darkness regardless of whether the fly still intends to walk in a fixed direction (Green et al. 2019). The trajectory shown in Figure 6 B indicates a consistent navigational goal throughout, even though the fly experienced three twelve-hour periods without a bar. In other words, each morning in which the bar reappeared after twelve hours of being off, it is as if the fly could pick up right where she left off in terms of traveling along her previous angle.

Figure 6 C shows the trajectories of all 29 flies tested in this paradigm, colored by fly. The twelve-hour nighttime periods with no bar are again plotted in black. These time periods appear as small spots with respect to the rest of the trajectory (again, because flies could not displace far without the bar). Qualitative inspection of these data suggests that the directions in which the flies walked before and after bar-off nights were largely similar. In other words, one could imagine removing the black “points”—which represent 12 hours of bar-off behavior—and not affecting the trajectories much.

The mean speed and tortuosity of flies tested in this intermittent-bar paradigm and the standard, constant-bar paradigm were similar (speed: 1.0 mm/s for each experiment, *p* = 0.86, tortuosity: 8.8 for the constant-bar flies vs 8.6 for intermittent-bar flies, *p* = 0.35, Mann-Whitney U test), suggesting that the general behavioral states of flies in the two experiments were comparable. For tortuosity, we excluded the bar off periods in the quantification. To quantify the difference in the fly’s goal angles across two daytime periods with an intervening night, we transformed each day’s trajectory into a vector from the start x-y location to the end x-y location on that day (Figure 6 D). We then calculated the angular difference between the vectors representing consecutive daytime periods, grouped by whether the intervening nighttime had the bar off or on (Figure 6 E). Independent of whether the bar was on or off in the intervening nighttime period, daytime pairs had a vector-angle difference that clustered strongly around 0 (Figure 6 F). The distribution of vector-difference angles for daytimes surrounding a bar-on night or a bar-off night did not show a statistically significant difference (*p* = 0.5, Mann-Whitney U test) (Figure 6 G).

To assess if the angular-difference measurements could have been observed by chance, we created a bootstrapped distribution of 10,000 mean absolute angular differences by randomly pairing days from all 29 flies, and calculating the mean absolute angular difference between each pair. If the headings were uniformly distributed, one would expect a mean absolute difference of 90^°^. Whereas if flies were to progress exactly along the same heading angle in two daytime epochs, one would expect a mean absolute difference of 0^°^. The measured mean absolute differences, surrounding bar-on (gray line, 23.9^°^) and bar-off (black line, 28.6^°^) nights fell well below the bootstrapped distribution (Figure 6 H), arguing that flies remember their angle even after 12 hours without the bar.

The observation that flies maintain a consistent direction after 12 hours of darkness suggests that one of the three previously mentioned possibilities is true: (1) the central complex may operate in a mode where the visual mapping to the EPG system remains stable for at least twelve hours, even in the absence of reinforcing visual input; (2) a fly performing long-term menotaxis may bypass the EPG visual-mapping process in generating this behavior; or (3) the EPG system may remap during prolonged darkness, but downstream circuits compensate, allowing the fly to maintain a stable allocentric heading despite the EPG phase changing its offset angle to the environment.

## Discussion

We built an open-source hardware and software system that enables routine virtual-reality behavioral experiments on many tethered *Drosophila* in parallel (Figure 1). The behavior of each individual can be studied at high spatiotemporal resolution for days to weeks. Flies on these rigs express naturalistic circadian and sleep rhythms (Figure 2), and they express a stable navigational goal direction on the timescale of days to weeks (Figures 3–6).

Our rigs resemble two other systems that allow for multiday behavioral neurophysiology in *Drosophila* (Huang et al. 2018; Flores-Valle et al. 2022; Flores-Valle et al. 2025). Although our platform is not currently optimized for simultaneous neural imaging, the small size and robustness of each individual rig allows for dozens of tethered flies to be studied in parallel for weeks at the behavioral level. Another published system allows for running visual neuroscience experiments in tethered *Drosophila* in a compact and economical manner (Loesche and Reiser 2021), but it does not currently allow for multi-day survival of individual flies. Our system’s hardware and software architecture is modular, allowing one to easily add new sensors and actuators or implement closed-loop real-time protocols that adapt experiments to the observed behavior on timescales of minutes to days. These rigs may thus enable a wide range of new experiments that were previously difficult to envision. For example, because our rigs allow for the delivery of thousands of sugar drops—which can act as behavioral reinforcers—over weeks, the rigs may ultimately help experimenters to operantly train flies to robustly perform trained tasks for the first time. Well-trained tasks are the bedrock of modern primate and rodent neuroscience. If individual *Drosophila* could be similarly trained, this would expand the range of questions that the field could ask and it may also help to better connect behavioral mechanisms discovered in mammals and flies down the road. This preparation may also enable scalable, high-resolution measurements of metabolic parameters in individual flies because it allows one to precisely control food intake over days while simultaneously monitoring behavior.

We discovered that flies exhibit a previously unrecognized ability to maintain a consistent goal angle over days to weeks, allowing them to displace tens to hundreds of meters from a start location. This behavior, which we call long-term menotaxis, involves flies progressing forward in a slower and more tortuous fashion than in short-term menotaxis (Green et al. 2019). Interestingly, when hungry flies find a small morsel of food, they express highly tortuous search trajectories centered on the food, presumably looking for more nourishment nearby (Bell et al. 1985; Kim and Dickinson 2017; Corfas et al. 2019; Behbahani et al. 2021; Goldschmidt et al. 2024). Because flies performing long-term menotaxis have trajectories with a tortuosity that is intermediate between that of flies performing short-term menotaxis and those performing local search, one possibility is that flies performing long-term menotaxis were alternating, on a seconds or minutes timescale, between a dispersal/menotaxis behavior and some other, more local-search-like behavior. Because the full range of motivations that give rise to long-term menotaxis—from hunger to social isolation to other unknown drives—remain unclear, extensive future work will be needed to explain the observed trajectories.

Flies performing long-term menotaxis had many extended periods—sometimes entire days—where an experimenter would be hard pressed to infer from their trajectory that there even existed a long-term goal angle (Figure 3 A, arrows). This observation argues that the goal angle is stored in a manner that can remain latent to behavioral measurement for long periods. An extreme example of this point is that flies can spend twelve hours walking in circles when an orienting cue is lacking. When the cue is restored they will readily return to their prior goal direction. The overall set of observations reported in this paper set a few constraints on the nature of the long-term goal signal. First, the long-term goal signal can stably persist for at least two weeks. Second, the long-term goal signal can persist during extended periods in which the fly is not actively pursuing it (even when an orienting cue is available and the possibility to pursue it exists). Third, the long-term goal can persist for at least twelve hours in situations where orienting cues are absent. Understanding the neural basis for a goal signal with these properties will be an important future direction for the field.

In the prevailing models of short-term menotaxis, the goal angle is stored in the same reference frame as the fly’s internal sense of heading, generated by EPG cells (Mussells Pires et al. 2024; Westeinde et al. 2024). If the longterm goal angle is also stored in reference to the EPG signal, then the fly’s ability to return to that goal angle after 12 hours without the bar implies that the offset between the EPG phase and bar position on the screen remains stable through twelve hours of complete darkness. However, in neurophysiological experiments that last approximately half an hour, experimenters already observe occasional, spontaneous changes in the offset between the EPG phase and the bar position on the screen (Seelig and Jayaraman 2015; Green et al. 2017). Thus, stability across twelve hours seems unlikely at face value. It is possible that the EPG-phase-to-bar-position map is more stable in our long-term experiments because the flies get several days of visual experience before the bar is turned off for twelve hours. This days-long experience might yield a visual map with a days-long persistence time, which would be remarkable. Alternatively, it is possible that the EPG phase does indeed remap often across twelve hours without a visual cue. In this case, the goal signal would need to somehow remap along with it. It is also possible that longterm menotaxis relies on some other, as-of-yet undescribed compass signal in the central complex that is more stable in its mapping to the visual world than the EPG bump. Finally, longterm menotaxis may not even rely on angular signaling in the central complex. Instead, it might be implemented in a manner that is completely different from short-term menotaxis. Whereas complete silencing of the EPG compass signal for days to weeks has proven challenging in our hands to date, experiments of this sort, once optimized, should help to differentiate among these possibilities.

Evidence exists that individual flies can express lifelong biases, or individuality, in aspects of their locomotor behavior—like turning more to the left or to the right, on average—and these biases are thought to reflect stochasticity in brain development across individuals (Ayroles et al. 2015; Buchanan et al. 2015; Linneweber et al. 2020). Our rigs may allow future work on such biases or proceed at extremely high spatiotemporal resolution, while also enabling one to ask how such biases interact with other motivational factors and learned behaviors to impact the overall behavioral output of insects over time. These studies on individuality have suggested that subtle but important anatomical differences across flies might account for the behavioral differences. Likewise, one possibility is that each fly relies on a stochastic developmental process to create a stable anatomical bias in the central complex, which promotes an individualized goal angle above all others over the whole lifetime of that animal.

To date, tethered behavior in *Drosophila* has largely focused on the minutes to hours timescale. The technical developments reported here should allow *Drosophila* behavioral biologists to routinely study the behavior of a dozen or more tethered flies over days to weeks. Beyond the ability to probe circadian and sleep rhythms with higher resolution and the discovery of a long-lasting navigational goal signal, this technology could enable the discovery of new behavioral phenomena that are only evident when examining flies closely for extended periods of time. If so, the advanced genetic and physiological methods in *Drosophila* should allow one to discover commensurate brain mechanisms that operate on such long timescales. These mechanisms are likely to transcend just developmental stochasticity and their description could have a broad impact on the field of systems neuroscience.

## Acknowledgements

We thank members of the Maimon laboratory for many helpful discussions. We thank the Rockefeller University Precision Instrumentation Technologies facility for their support in building and testing various parts of the rigs, and for many helpful discussions. We thank Frank Loesche and Michael Reiser for generously sharing their progress on a ROS2 adaptation of Fictrac. Research reported in this publication was supported by a Brain Initiative grant from the National Institute of Neurological Disorders and Stroke (R01NS104934) to G.M and an R35 grant from the National Institute of Neurological Disorders and Stroke (R35NS132252) to G.M. G.M. is a Howard Hughes Medical Institute Investigator.

## Author Contributions

J.W. and G.M. conceived of the project. J.W. designed and built the experimental rigs, performed the experiments, and analyzed the data. M.W. collected data for Figure 3. J.W., T.M., and J.R. developed and implemented the software system for the experiments. E.D.-F. designed and tested the initial sugar-pipette feeding system in similar behavioral rigs, showing that head-fixed flies could survive for up to ∼2 weeks. J.W. and G.M. jointly interpreted the data and decided on new experiments. J.W. and G.M. wrote and edited the paper.

## Methods

### Experimental system overview

We developed experimental rigs, each of which allowed a single head-fixed fly to survive for up to two weeks within a visual virtual-reality environment. Individual flies walked on an air-supported ball and were surrounded by a panoramic screen, and were fed small sugar drops at regular intervals. A projector was used to display visual stimuli on the screen. Sugar-drop delivery and other electronic hardware, like LEDs, were regulated by a custom controller box. We next provide further details on each rig component.

### ROS system

To control experiments, generate visual stimuli, and integrate input and output signals, we developed a software environment using the Robot Operating System 2 (ROS2) libraries. A single Linux computer controls each set of three rigs. ROS2 allows fast and flexible communication between many software processes, including those driving hardware. ROS2 provides a language-agnostic middleware for code written in multiple programming languages. This feature allows performance-critical processes, like the ball tracking software to be written in a compiled language (e.g., C++), while other aspects of the system, such as the experiment control code, to use an interpreted language (e.g. Python).

A functional unit in a ROS2 environment is called a node, with different nodes communicating with each other via custom ROS messages, asynchronously, during an experiment. This distributed architecture facilitates an experimental setup where core nodes are written, tested, and rarely changed, and thus only requires the experimenter to customize a few experiment-control nodes in order to implement new experiments. A flexible Graphical User Interface (GUI) displays current information about the experiment and allows for real-time user interaction with the rig. Fictrac, the ball pose tracking software, has been modified to fit within this ROS2 environment.

Each experiment is defined by one or more YAML configuration files, that define which nodes to launch, as well as the parameters to be used by each node. A custom “bringup” package parses the configuration files, and launches the appropriate ROS2 nodes with the specified parameters. This program also manages the output data directory structure, and distributes this information to each node so that they may save their data in an appropriate location. A copy of the configuration file, as well as the current development state of each of node to be launched, is saved in the output folder, so that the state of the experiment can be exactly reproduced at a later date. Overall, this system allows for quick and convenient adjustments to experimental design parameters via edits to the configuration files. A graph of the information flow between main components of the rig, and the ROS2 nodes that interact with them is shown in Figure S1 D.

### Visual Display System

In order to form a panoramic image around the fly, we back-projected onto a truncated cone (a frustum) Figure 1 C, using a portable projector (Texas Instruments Lightcrafter 3010) (Creamer et al. 2019; Longden et al. 2023). Running in its standard mode, this projector has a flicker rate of 240 Hz, well above *Drosophila*’s flicker fusion frequency (Miall 1978) and a maximum frame rate of 120 Hz. The standard version of the projector employs LEDs with center wavelengths of 623 nm, 540 nm, and 456 nm, but these are easily swappable if other wavelengths are desired (Longden et al. 2023).

Unlike past approaches for projector-based panoramas in *Drosophila* (Haberkern et al. 2019; Creamer et al. 2019), we adapted this approach from the rodent literature (Aronov and Tank 2014) because it requires only one projector. For steric reasons, we reflected the projected image off a 45^°^ first surface mirror (firstsurfacemirror.com) (Figure 1 C). The screen used herein displayed an image that covered 320^°^ around the azimuth and from 10^°^ above the fly’s horizon to 50^°^ below it. These angles were chosen because they encompass the full field of view afforded to a fly affixed to the standard pyramid-shaped stage we use in behavioral neurophysiology experiments (Maimon et al. 2010). Screens with larger or smaller fields of view are easily accommodated.

The frustum screen was 1.25 in high, with a minor radius of 0.75 in and a major radius of 2 in (Figure S1 A,B). For the behavioral experiments reported on here, the screen was made out of 80-lb cardstock (Desktop Publishing Services, Part 59421–50). The screen had a gap of 40^°^ in azimuth where no image was projected. This gap is roughly the size of the blind spot of *D. melanogaster* (Buchner 1971; Heisenberg and Wolf 1984; Zhao et al. 2025). The visual stimuli were rendered by the Projector Driver ROS Node and sent to the projector (Texas Instruments LightCrafter DLP3010EVM-LC) at 120 Hz over HDMI. (Though due to computation time on the software end, frame updates were often slower than 120 Hz.) Projected images were properly warped to compensate for the geometry of the projected image and screen. The final, relative positions of the projector, mirror, and conical screen were adjusted for each experimental setup by hand—via a calibration image of concentric rings—to account for slight variability in the optical axis of different projectors, despite the projectors being nominally identical.

### Rig controller box

A custom controller box, organized around a printed circuit board that includes an Arduino-compatible microcontroller (Adafruit Metro Mini), allowed the computer (via USB communication) or a human user (via knobs and switches) to regulate the rig’s hardware (Figure S1 C). The controller box drove 240, white, low-power LEDs that surrounded the top perimeter of the enclosure. These LEDs generated overhead lighting that turned on and off at regular intervals (12 hours on, 12 hours off), providing flies with a circadian entrainment cue. The controller box could also drive up to four high-power LEDs and two diode lasers. In the experiments reported here, we used two high-power infrared (850 nm) LEDs to illuminate the fly and the floating ball (via fiber-optic light guides). We did not employ any diode lasers; these can be used to rapidly heat up the fly above room temperature to motivate locomotion or as an aversive cue for associative learning. For the feeding servo motor, a button could induce the pipette to toggle between the inward feeding position and the outward retracted position, which was necessary for calibrating the precise feeding position. There is also a button that manually triggered the solenoid to release a sugar drop and a rotary encoder knob adjusted the open time of the solenoid. An LCD display indicated the solenoid open time and IR laser intensity levels. The schematic of the controller’s PCB is shown in Figure S2, broken up into its main functional blocks. The controller box also drove a solenoid to enable the feeding system described below.

### Feeding Device

To keep flies alive, we fed them drops of a sucrose solution, similar to Götz (1987). Richer diets, which contain amino acids and other nutrients, could be used in the future (Lee et al. 2008; Ja et al. 2007; Yapici et al. 2016). To deliver sucrose drops to the fly, the feeding device moved a pulled glass capillary, or pipette, containing the sucrose solution from a position ∼4 mm away from the fly to just under the fly’s proboscis. Once the pipette was under the fly’s proboscis, we ejected a ∼15 nL drop of ∼150 mM (5% w/v, as in (Ja et al. 2007)) sucrose solution, via a solenoid valve (adapted from Forman et al. (2017)), which the fly consumed. After consumption, the pipette returned to its rest position 4 mm away, which prevented the fly from interacting with it between feeding events.

The feeding device consisted of a linear servo motor (Actuonix L16–50–35–06-R), mounted in a custom 3D-printed holder and connected via another 3D-printed part to a pipette holder (5430-ALL, World Precision Instruments), which held a pulled glass capillary (pipette) (Figure 1 B). The two 3D-printed parts connect this assembly to a linear ball-bearing guide rail (MGN7 C, 100mm, AliExpress), which constrains its motion to a single axis. This increased the precision of the positioning of the pipette to ∼33 *µ*m (s.d) on the axis of motion and ∼5 *µ*m (s.d.) on the perpendicular axis. Feeding pipettes are pulled from borosilicate glass capillaries (Sutter Instrument BF150-110-7.5) using a micropipette puller (Sutter Instrument P-1000). The pipettes were stabbed through a stretched Kimwipe to give a beveled opening of approximately ∼30 µm in diameter. Pipettes were backfilled with 5% sucrose solution and flies were fed drops of this solution, which emerged from the tip via pressurized pulses of air.

Drop volume was adjusted for each experiment manually, using the controller box to change the open time of the solenoid valve that delivered the pressure pulse. The air pressure was set to 3 psi using a precision pressure regulator (Festo LRP-1/4–4), and the solenoid open time was set to a duration of ∼20 ms-80 ms.

### Behavioral imaging

The ball camera (FLIR Chameleon3 CM3-U3-13Y3M-CS) was fitted with an Infinistix 0.45x 94mm WD lens (Infinity photo optical, 194045) and an 850 nm, 50 nm bandpass filter (Edmund Optics, 84-778), held in place in the back of the lens by a 3D-printed adaptor. Ball tracking frames were collected and analyzed by Fictrac at 60 Hz. Another camera (FLIR Chameleon3 CM3-U3-13Y3M-CS) viewed the fly from the side via a variable focus lens (Computar MLM3X-MP). This camera ran at 30 Hz for monitoring the fly in real time. To avoid excessively large videos, images from this camera were only saved at 0.6 Hz. We also saved a 5 s burst of frames, at 30 Hz, triggered by the beginning of each sugar-drop feeding event, which could be used to verify if the feeding took place as expected.

### Physical construction

Each rig was assembled on one third of the footprint of a 24 ×16 in breadboard (Thorlabs B2448FX or Thorlabs MB2448). We constructed a frame around each rig, consisting of 1 inch t-slotted aluminum rails (McMaster Carr 47065T503) with walls made out of 1/4 in thick expanded PVC foam board (US plastics 44170). This frame provided mechanical stability and light isolation. The the second set of three rigs mounted onto the frame of the bottom set.

### Fly husbandry

Canton S flies were reared in standard cornmeal-agar-molasses food, supplemented with 100 µg/ ml ampicillin, in incubators kept at 25^°^C. Ambient lighting turned on and off with a twelve-hour cycle. In the timed feeding experiments (Figure S7), flies were reared in vials where ampicillin was not included in the food.

### Experimental protocols

Flies were tethered via blue-light activated glue (Bondic) to 0.005 inch tungsten pins two to three days post-eclosion. They were anesthetized by chilling to 3–4^°^C on a Peltier cooler (Teca LHP-300CP) driven in closed loop by a PID controller (Teca TC-3400). The wings were glued together with a small drop to prevent flight attempts while on the ball. Flies were kept in a small chamber humidified with a wet Kimwipe to recover between tethering and being placed on the rigs.

The 6.35 mm diameter ball that the fly walked on was carved by hand from Last-A-Foam FR-4618 (General Plastics) as previously described (Green et al. 2017). The ball sat in a machined aluminum holder, and was supported on a cushion of air flowing at ∼ 0.5 L/min, as previously described (Green et al. 2017). This air was humidified by passing through the headspace of a ~ 4 L jar filled 20 – 80 % with DI water. Humidfying the ball’s airflow seemed to aid fly survival across days.

Flies were not explicitly heated in the longterm experiments, and they experienced a temperature between ∼24 – 26 ^°^C given the interaction of the room’s air conditioning and the heat generated by the electronics in the rig. Flies were fed according to the rule described in the main text, with drops of 5 % (w/v) sucrose solution.

For short-term menotaxis experiments (Figure 3 B, D, F), flies were explicitly heated by elevating the temperature of the air that was used to float the ball. The ball air was heated via current that was passed through nine inches of nichrome wire (McMaster Carr, 8880K77) that was wrapped around the aluminum ball holder. The nichrome wire was electrically insulated from the aluminum ball holder by a layer of polyimide tape (McMaster Carr, 1754N12). A thermistor was affixed to the aluminum ball holder to monitor its temperature (Digikey, 495–2149-ND). The thermistor, aluminum ball holder, and nichrome wire were adhered together, to form a single thermal unit, via thermally conductive epoxy (MG Chemicals, 8329TCM). A second custom controller box maintained the ball holder at a consistent target temperature using a PID control loop. This temperature was monitored and controlled by the ROS2 software system. For the experiments described here, the ball holder was set to 38 ^°^C, leading to a temperature of ∼33.5 ^°^C around the fly.

## Data Analysis

Each node in the ROS system described here may produce and save data as separate files. All time series data were saved in the common timebase of the computer clock. Data were analyzed using custom Python code. Raw Fictrac data from a single fly, collected for one week at 60 Hz, produced ∼36 million timepoints and a file size of ∼20 GB. For most analyses, we downsampled the trajectory data to bins ranging from 1 s to 30 min. The bin size was selected based on the level of detail required for each analysis. Specifically, for the main experiments in Figures 2–4 we used a 1 s time base everywhere except where otherwise noted, and for the trajectories shown in Figure 3 A, E, we used 10 s bins. For the virtual rotation experiment in Figure 5, we also used 10 s bins, except in the trajectories close to two bar jumps, (Figure 5 B), where we used the full 60 Hz data. For the intermittent-bar experiment in Figure 6, we also used 10 s bins.

A fly’s traveling speed for each 1 s time bin was calculated by taking a vector sum of the forward and sideways translations within the 1 s bin. Speeds in longer time bins were calculated by averaging the speeds calculated in the 1 s bins.

To calculate tortuosity and circular variance of heading metrics for the trajectories of flies performing short-term menotaxis (Figure 3 B, D, F), we calculated both metrics for each 30 min short-term menotaxis bout that each fly conducted and used the mean across the two bouts. For the circular variance comparison shown in Figure S6 A, we excluded any timepoints where the fly was standing still (speed < 0.25 mm/s), and compared the distributions of circular variances of heading angles between heated and unheated flies using a Mann-Whitney U test.

### Circadian Time Compensation analysis

To test whether the flies were compensating their travel direction for the circadian movement of the bar/sun, we rotated each fly’s trajectory by the azimuthal angle of the sun throughout each the day, and tested whether these trajectories were more or less tortuous after this adjustment. Because it is unclear what the prediction should be at night, we restricted this analysis to daytime periods, concatenating the trajectories from all days, and then rotating them by the azimuthal angle of the sun at each timepoint.

The way the sun moves in the sky is called the solar ephemeris function. This function differs across different points on the earth, and during different seasons. The Solar ephemeris function describes the differences in the sun’s movement across the sky due to latitudinal and seasonal variations. Since the flies always experienced 12 hours of daytime from the overhead lights, we only used the solar ephemeris function accurate during either equinox, when the sun is up for 12 hours, thus matching the flies’ experiences, as shown in Figure S9 A. We calculated the solar ephemeris on the equinox for 9 latitudes from-80^°^to 80^°^, and rotated the trajectories accordingly. Sunrise and sunset were aligned to the times when the overhead lights turned on and off, respectively.

An example fly’s trajectory before and after these rotations is shown in Figure S9 B. The tortuosity for each modified trajectory, compared to the original trajectory, is shown in Figure S9 C. For each fly, the lowest tortuosity achieved by any of the nine solar-rotation manipulations was compared to the original tortuosity, as shown in Figure S9 D. Tortuosities are significantly higher after solar-rotation manipulations (Wilcoxon signed-rank test, *p* = 1.3 × 10^−5^). All but one fly showed an increase in tortuosity after all solar-rotation manipulations.

## Code and data availability

All data, code, and detailed electronic and CAD designs associated with these systems will be made publicly available upon publication. Requests pre-publication can be directed to the authors.

## Supplemental Figures

**Figure S1:**
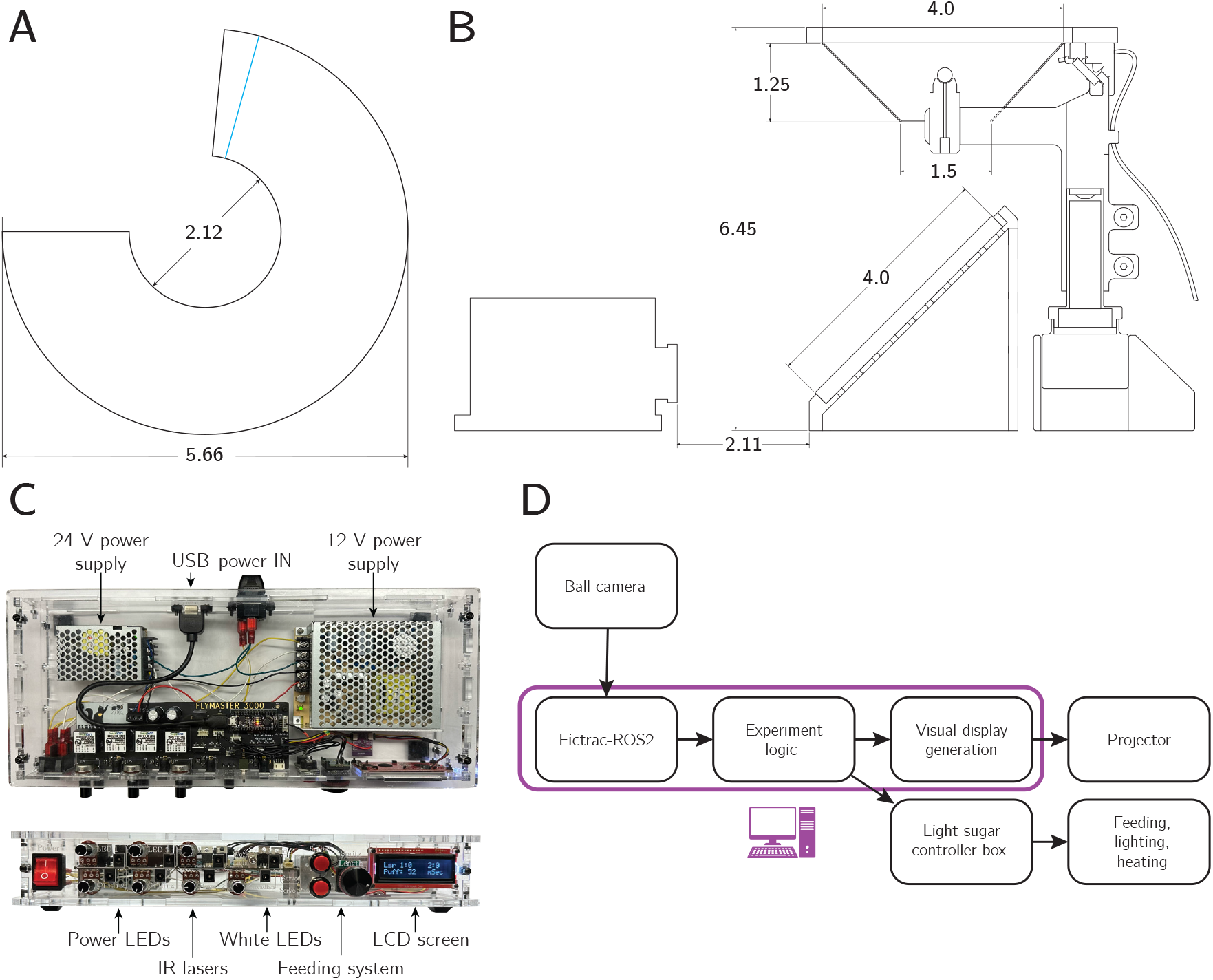
**A**: The template for cutting out the conical screen. The blue line marks where the opposite edge should line up to form the correct frustum shape, and can be engraved with the laser cutter to mark it. All dimensions are in inches. **B**: Dimensional drawing of the display assembly. **C**: Top and front views of a controller box that drives electronic components on the rig. **D**: Schematic of information flow through the rig

**Figure S2:**
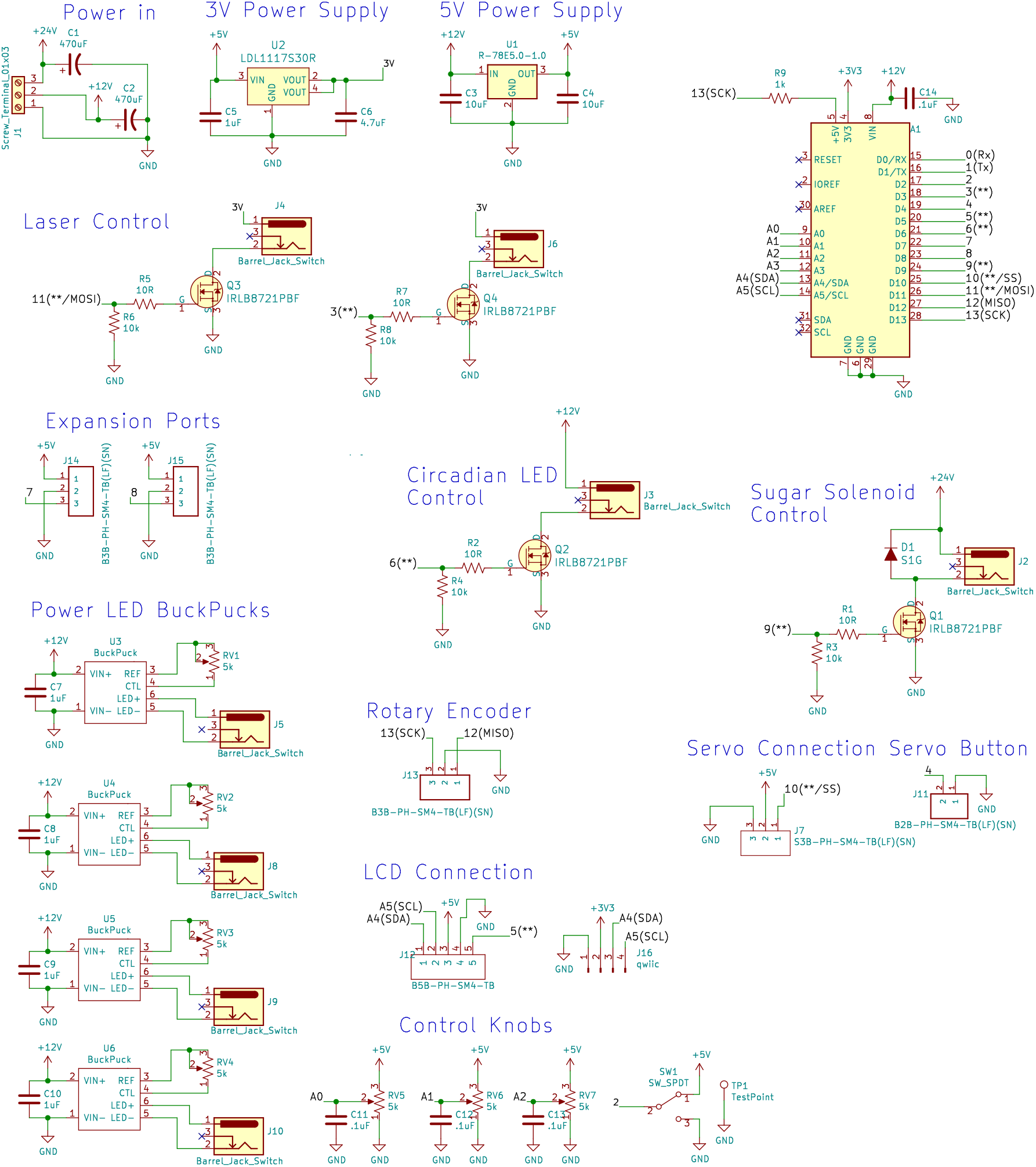
Schematic for the controller box PCB.

**Figure S3:**
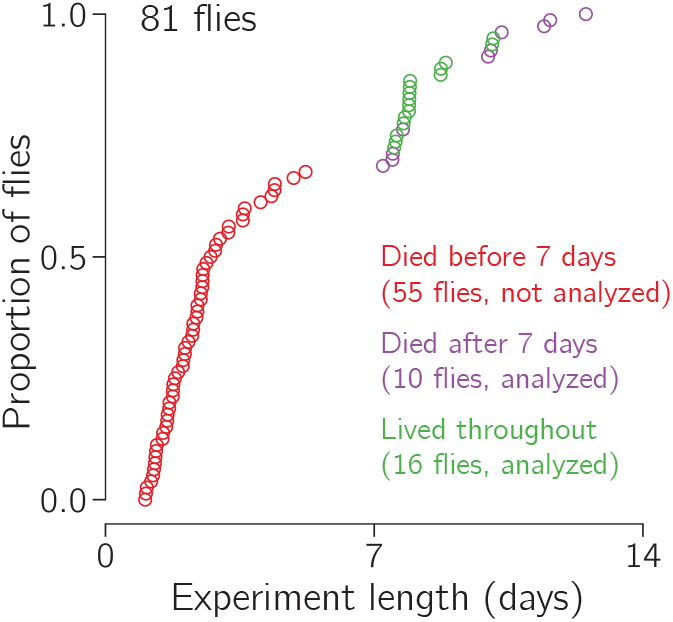
Survival curve for the flies analyzed in Figures 2 and 3. Each fly’s survival time is indicated by a circle. Red circles indicate flies that did not live long enough to be included. Purple circles indicate flies were included but died while on the rigs. Green circles indicate flies that were removed from the rigs without dying.

**Figure S4:**
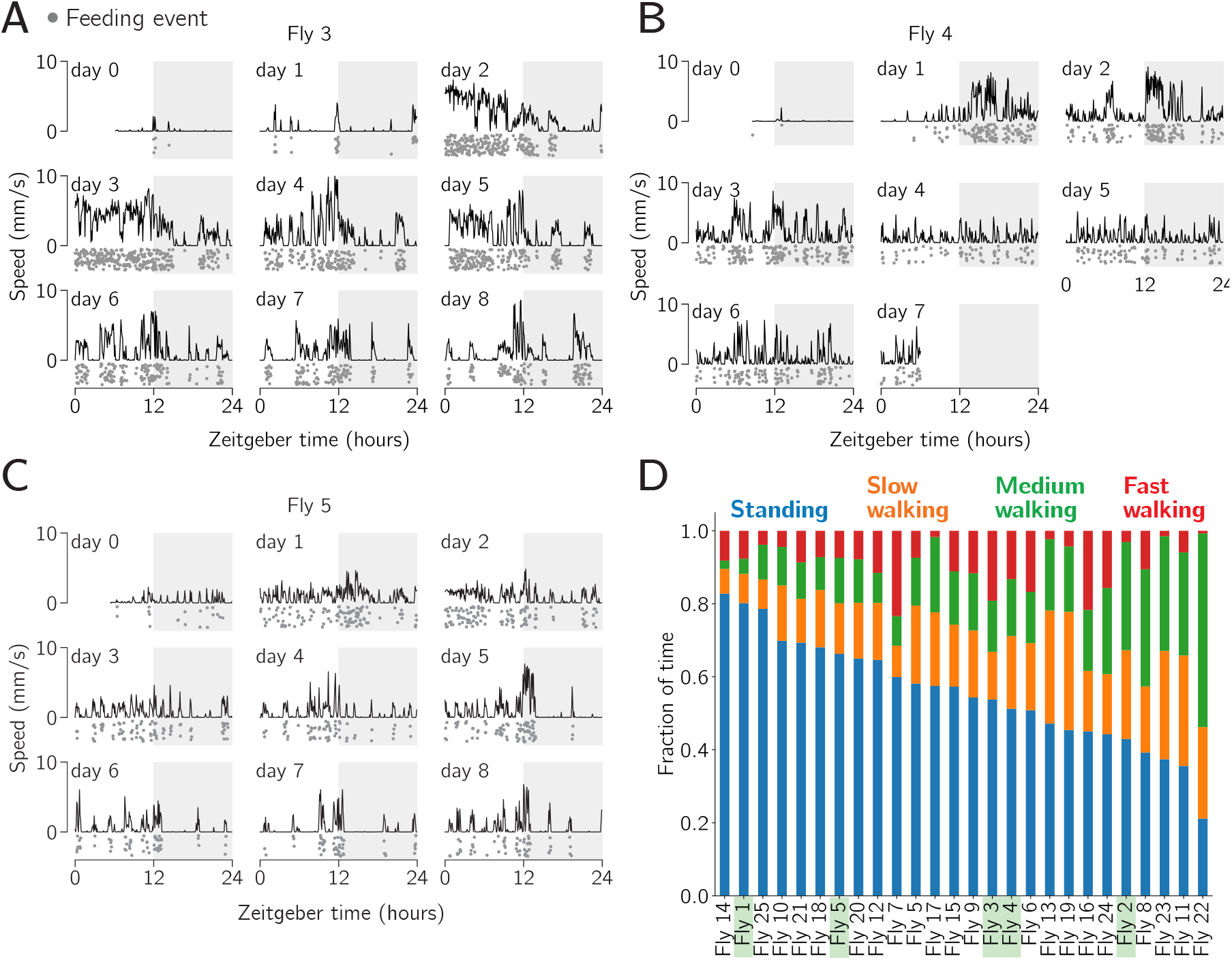
Individual flies show different walking dynamics. **A-C:** Translational speed over time for three example flies (1 min bins). Each feeding event is indicated by a gray dot below the speed trace (each dot’s *y* position is jittered for clarity). **D:** Binned translational speed profiles for all 26 flies in the data set. For each fly, the mean translational speed was calculated in each minute and assigned to a behavioral bin as follows: standing (<0.1 mm/s), slow (0.1 – 1 mm/s), medium (1 – 3 mm/s), or fast (*>*3 mm/s). Flies are sorted by the standing fraction. The example flies from panels A-C, Figure 2 C-D, and Figure 3 B and D, are highlighted in green.

**Figure S5:**
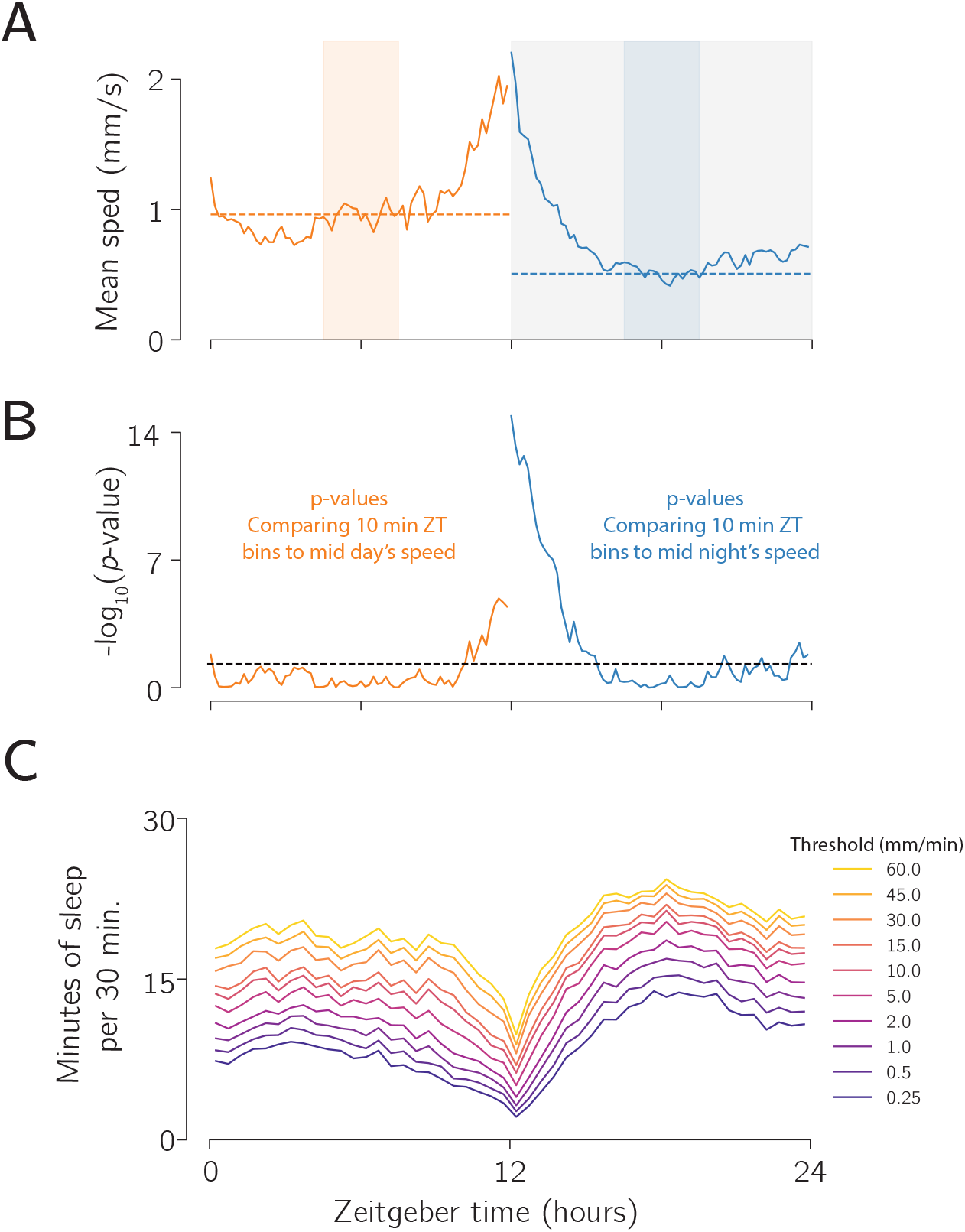
Flies walk significantly faster in anticipation of the lights turning on and off. **A:** Mean speed across all flies as a function of Zeitgeber time. The middle three hours of the day (orange shaded region) and night (blue region) form the comparison group for the rest of the day / night timepoints. The mean speed for each is comparison group is shown as a dotted line. **B:** Plot of the log_10_ of the *p* value for the comparison of the distribution of mean speeds for each 10 minute bin compared with the mid-day or mid-night time window (Mann-Whitney U test). Significance threshold of *p* = − 0.05 (not corrected for multiple comparisons) is shown as a dashed line. **C:** Average sleep per 30 minute bin parsed by the speed threshold used for defining a standing event.

**Figure S6:**
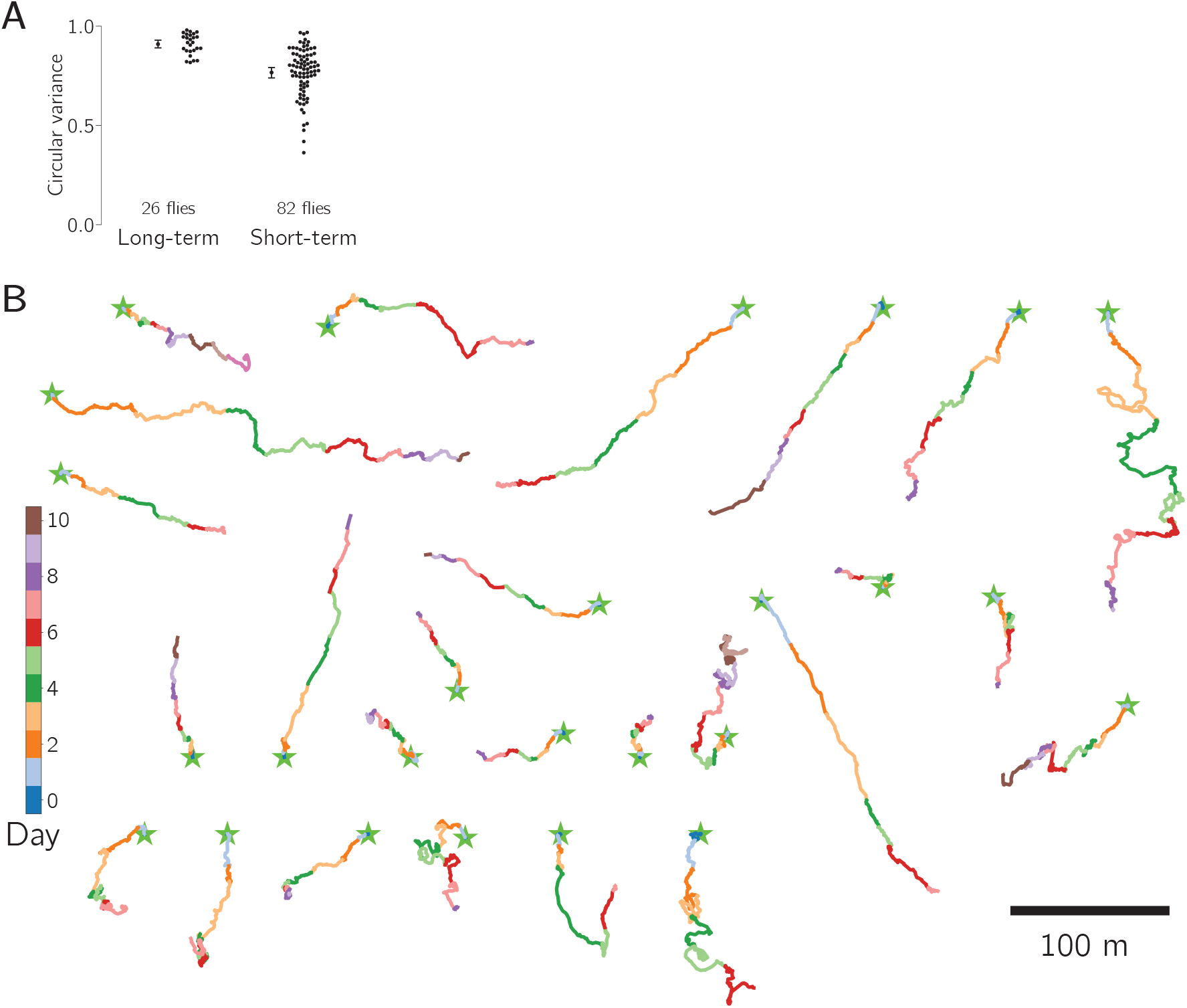
Circular variance analysis and separated trajectories of long-term menotaxis behavior, relating to Figure 3. **A:** Flies performing long-term menotaxis show significantly higher circular variance in their headings angle while walking compared to flies performing short-term menotaxis. **B:** All trajectories from Figure 3 F, shown separated and colored by day of the experiment. Trajectory start locations are shown as green stars.

**Figure S7:**
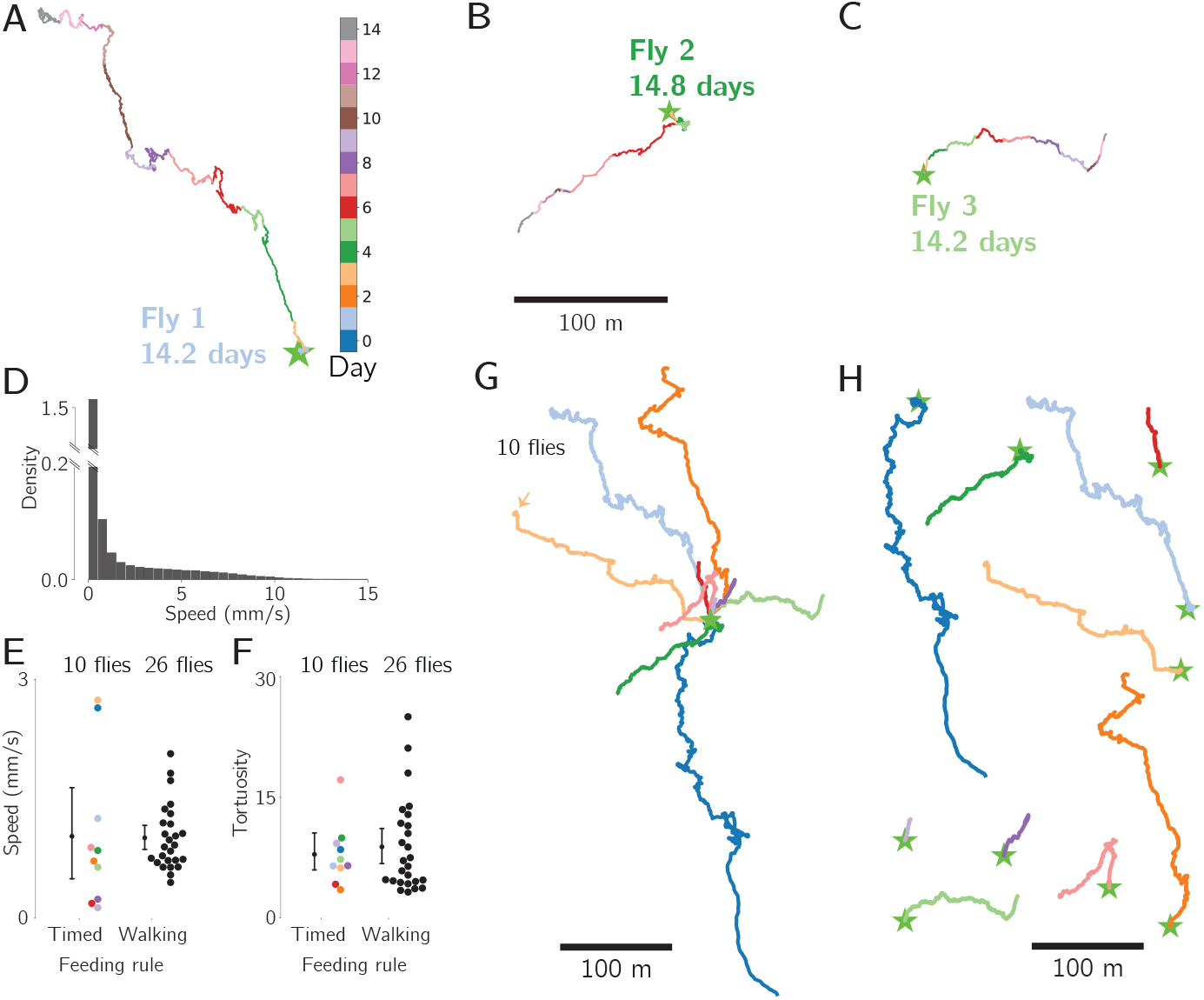
Flies fed every 15 min expressed a preferred traveling direction for up to two weeks. **A, B, C:** 14-day trajectories from three flies. These flies were fed a sugar drop every 15 min. The start of each trajectory is indicated with a green star in all panels. **D:** Histogram of average walking speed calculated in 1 s bins for all flies (standing events excluded). Bin width: 0.5 mm/s. Note discontinuous axis on the first bin. **E:** Comparison of average speed per fly from this data set (timed feeding) with the long-term menotaxis flies from Figure 3 D (walking-triggered feeding). The average speed per fly was not significantly different (Mann-Whitney U test, *p* = 0.4). Mean and bootstrapped 95% confidence intervals are shown. **F:** Tortuosity for each fly’s trajectory from this data set (timed feeding) and the flies performing long-term menotaxis flies from Figure 3 D (walking-triggered feeding). The tortuosity per fly was not significantly different between the two groups (Mann-Whitney U test, *p* = 0.99). **G:** Trajectories of all flies from this data set. Individual fly colors match those in panels E, F, and H. All flies were tested for 10–15 days, except for the fly plotted in light orange, and indicated with an arrow, which was tested for just over 5 days. **H:** Same trajectories as in panel G, but shown displaced from one another.

**Figure S8:**
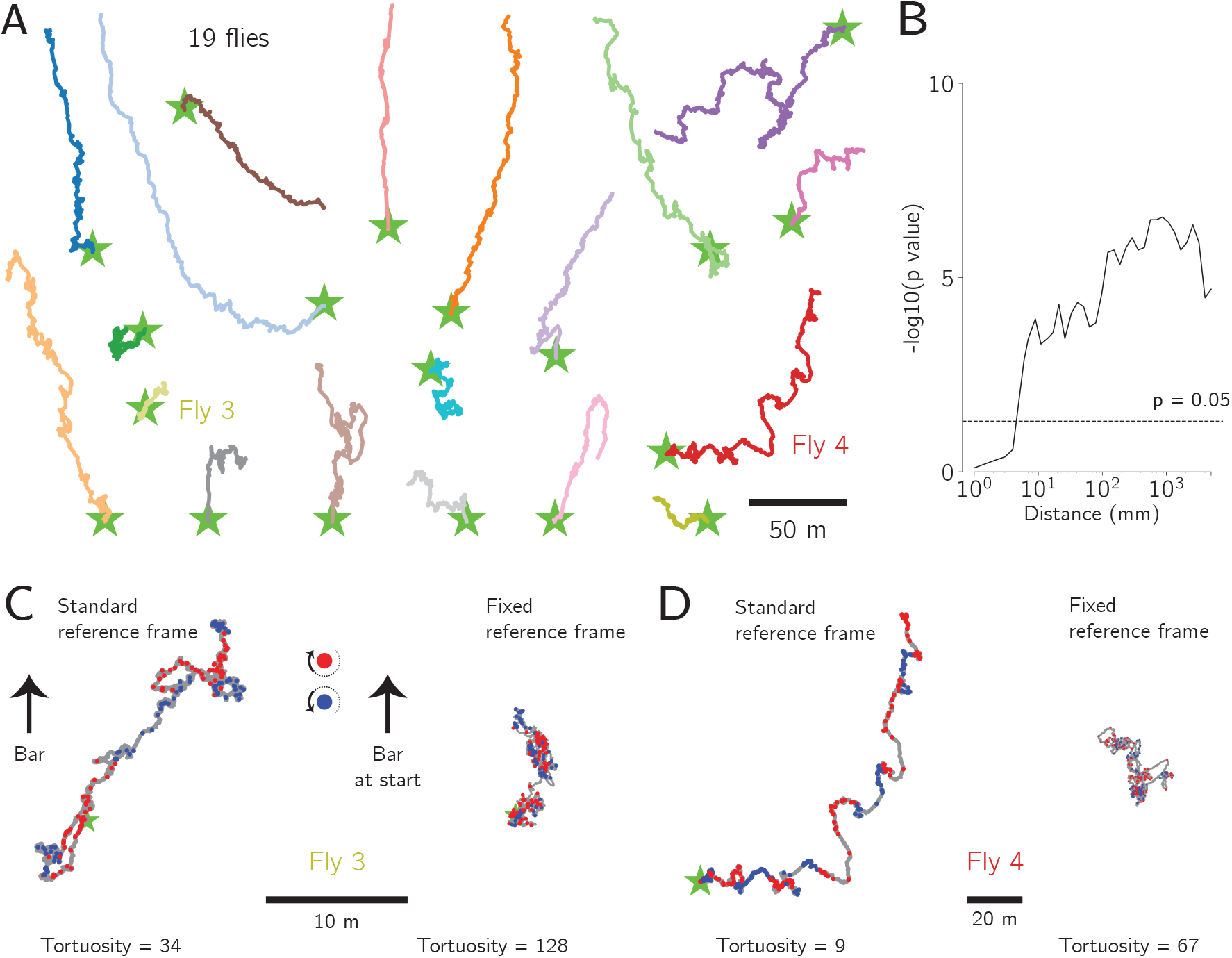
Additional analyses of the virtual rotation experiment. **A:** Entire trajectories for all 19 flies tested in the virtual rotation experiment of Figure 5. Colors associated with individual flies here match those used in Figure 5 E and F. **B:** Plot of the log_10_(*p*) value vs path distance for the analysis described in Figure 5 C. The dashed line indicates the *p* = −0.05 significance threshold (not corrected for multiple comparisons). **C:** Comparison of the trajectory of an additional example fly in the standard vs fixed reference frame plots of Figure 5. Despite the fact that this example fly had a more tortuous trajectory in the standard frame (tortuosity 34), her trajectory was still even more tortuous in the fixed frame (128). **D:** Same as panel C, for one more fly.

**Figure S9:**
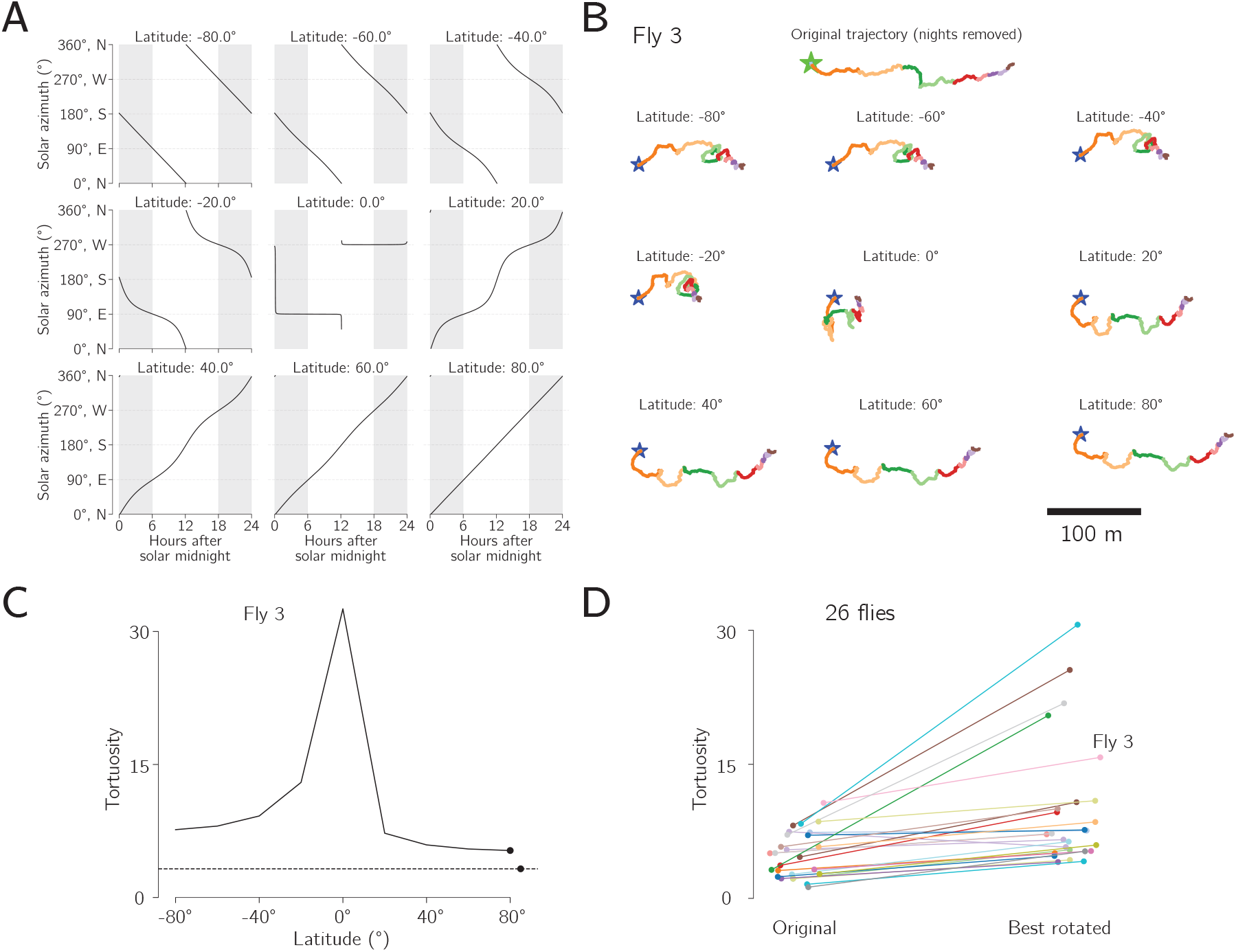
No evidence for circadian time compensation in long-term menotaxis. **A:** Solar ephemeris functions for nine different latitudes, calculated for the equinox. Each plot shows the solar azimuthal angle (degrees) as a function of time after solar midnight. Night time is shaded in gray. **B:** One example fly’s trajectory with nights removed (original, top), and the same trajectory rotated according to the solar ephemeris function for each of the nine latitudes (bottom nine plots), labeled by latitude. **C:** Tortuosity of the fly from panel B’s trajectory (original, dashed line) compared to the tortuosities of the nine solar-rotated trajectories. **D:** Comparison of the tortuosity of all flies original trajectories vs the lowest tortuosity achieved by rotating according to any of the nine solar ephemeris functions. Tortuosities are significantly higher after solar-rotation manipulations (Wilcoxon signed-rank test, *p* = 1.3 ×10^−5^). All but one fly showed an increase in tortuosity after all solar-rotation manipulations.

## Notes

### Competing Interest Statement

The authors have declared no competing interest.

